# The genomic architecture of competitive response of *Arabidopsis thaliana* is highly flexible between monospecific and plurispecific neighborhoods

**DOI:** 10.1101/536953

**Authors:** Cyril Libourel, Etienne Baron, Juliana Lenglet, Laurent Amsellem, Dominique Roby, Fabrice Roux

## Abstract

Although plants simultaneously interact with multiple neighboring species throughout their life cycle, there is still very limited information about the genetics of the competitive response in the context of plurispecific interactions. Using a local mapping population of *Arabidopsis thaliana*, we set up a Genome Wide Association study to estimate the extent of genetic variation of the competitive response in presence of 12 plant species assemblages, and to compare the genetic architecture of the competitive response between monospecific and plurispecific neighborhoods. Based on four phenotypic traits, we detected strong crossing reaction norms not only among the three monospecific neighborhoods, but also among the different plant assemblages. Accordingly, the genetic architecture of the competitive response was highly dependent on the identity and the relative abundance of the neighboring species. In addition, enriched biological processes underlying the competitive response largely differ between monospecific and plurispecific neighborhoods. In particular, receptor-like kinases and transporters were significantly enriched in plurispecific neighborhoods. Our results suggest that plants can integrate and respond to different species assemblages depending on the identity and number of each neighboring species, through a large range of genes associated mainly with perception and signaling processes leading to developmental and stress responses.

## Introduction

Because plant-plant interactions are recognized as a major factor mediating plant community structure, diversity and dynamics (Tilman 1985; Goldberg & Barton 1992; Chesson 2000; Martorell & Freckelton 2014), deciphering the genetic and molecular bases underlying plant-plant interactions appears fundamental to predict the evolutionary dynamics of plant communities in ecological time (Pierik et al., 2013; Frachon et al., 2017). This is especially relevant in the context of current anthropogenic modifications of plant assemblages, which may in part result from the intertwined effect of increased plant biomass and reduced plant diversity under climate warming (Baldwin et al., 2014) or from native species having different geographical range shifts under climate change (Bachelet et al., 2001; Gilman et al., 2010; Singer et al., 2013). Because the average potential to reduce crop yield is significantly higher for weeds than any crop pests (Oerke et al., 2004; Neve et al., 2009), identifying and characterizing the function of genes underlying plant-plant interactions appears also fundamental to accelerate breeding programs aimed at increasing crop competitiveness (Worthington & Reberg-Horton 2013; Onishi et al., 2018). In addition, in the context of complementarity in using resources, optimizing species assemblages in cropping species may be facilitated by the understanding of the genetics underlying overyielding (Litrico & Violle 2015; Pakeman et al., 2015; Weiner et al., 2017).

However, in comparison to other types of biotic interactions such as plant response to virus, bacteria, fungi, oomycetes and to a lesser extent herbivores, there is still very limited information about the genetics associated with natural variation of plant-plant interactions, i.e. when plants have been directly challenged by other plants (and not in artificial environments simulating plant-plant interactions) (Bergelson & Roux 2010; Bartoli & Roux 2017). For example, a recent review listed only 47 Quantitative Trait Loci (QTL) mapping studies (including three Genome Wide Association studies, GWAS) that have been designed to study the genetic architecture underlying natural variation of plant-plant interactions (Subrahmaniam et al., 2018). About two-thirds of these QTL mapping studies focused on asymmetric interactions (i.e. when one of the interacting partners benefits at the expense of the other), including allelopathy underlying weed suppressive ability and response to parasitic plants (Subrahmaniam et al., 2018). Surprisingly, despite the importance of competition in driving plant community assemblages, only six QTL mapping studies (including two GWAS) focused on competitive interactions in a heterospecific context, i.e. when both interacting species suffer significant cost by investing in competing and therefore compromising on the benefit (Dudley 2015). In agreement with other types of biotic interactions (Roux & Bergelson 2016; Bartoli & Roux 2017), plant-plant interactions are mainly driven by a complex genetic architecture, ranging from the identification of few medium-effect QTLs to the identification of up to tens of small-effect QTLs (Subrahmaniam et al., 2018).

While informative, most of these QTL mapping studies are based on monospecific heterospecific interactions (i.e. one single pair of interacting species; Subrahmaniam et al., 2018). However, throughout their life cycle, focal plants often interact simultaneously with several neighboring species, either in crop fields or in natural communities (Wilson et al., 2012). This highlights the need to study the genetic architecture underlying plant-plant interactions by considering the response of a focal species to plurispecific interactions. In particular, whether the genetic architecture underlying the response of a focal species in a plurispecific neighborhood corresponds to the sum of QTLs that are specific to a neighbor species and/or to the emergence of new QTLs remains on open question (Subrahmaniam et al., 2018).

To address this question, we set up a GWAS to compare the genetic architecture of competitive response of the model plant *Arabidopsis thaliana* between monospecific and plurispecific neighborhoods. Although *A. thaliana* has long been considered as not being often challenged by other species in natural plant communities, several studies recently challenged this view (i) by revealing extensive genetic diversity associated with the response to interspecific competition (Brachi et al., 2012; Bartheleimer et al., 2015), in particular at the within-population scale (Baron et al., 2015; Frachon et al., 2017), and (ii) by finding an adaptation of a genetically polymorphic local population likely to increased interspecific competition in less than eight generations (Frachon et al., 2017). In addition, in the context of monospecific interactions, the genetic architecture of competitive response of *A. thaliana* was found to be highly dependent on the identity of the neigboring species (Baron et al., 2015), making *A. thaliana* an attractive model to test the absence or presence of new QTLs in a plurispecific neighborhood in comparison to related monospecific neighborhoods.

In this study, we first estimated the extent of genetic variation of competitive response in a set of 96 local *A. thaliana* French accessions, which were submitted to monospecific and plurispecific competition treatments based on all one-way, two-way and three-way combinations of three species frequently associated with *A. thaliana* in natural plant communities in France, *i.e. Poa annua, Stellaria media* and *Veronica arvensis*. Based on the whole-genome sequence of the 96 accessions, we then run GWA mapping to compare the genetic architecture of competitive response of *A. thaliana* between monospecific and plurispecific neighborhoods. Finally, we examined whether biological processes overrepresented among SNPs involved in competitive response were different between monospecific and plurispecific neighborhoods and discussed the function of candidate genes.

## Materials and Methods

### Plant material

A set of 96 whole-genome sequenced accessions of *A. thaliana* collected in the TOU-A local population (Toulon-sur-Arroux, Burgundy, France, 46°38’53.80″N - 4° 7’22.65″E) were used for the purpose of this study. As previously described in Frachon et al. (2017), the TOU-A population is highly polymorphic at both the phenotypic and genomic levels. Importantly, the very short Linkage Disequilibrium (LD) observed in this population 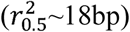 allows the fine mapping of genomic regions associated with natural variation of phenotypic traits down to the gene level (Brachi et al., 2013; Huard-Chauveau et al., 2013; Frachon et al., 2017).

Maternal effects of the 96 accessions were reduced by growing one plant of each family for one generation under controlled greenhouse conditions (16-h photoperiod, 20°C) in early 2011 at the University of Lille. Given an estimated selfing rate of ∼94% in this population (Platt et al., 2010), the 96 accessions were considered as mostly homozygous along the genome.

In this study, we used three neighboring species commonly associated with *A. thaliana* in natural plant communities in France and detected in the TOU-A plant community (F. Roux, personal observation). These species are the meadow grass *P. annua* (Poaceae) with a low spreading growth form, the chickweed *S. media* (Caryophyllaceae) and the speedwell *V. arvensis* (Scrophulariaceae) both with a crawling growth form. Seeds for these three species have been obtained from the Herbiseeds company (http://www.herbiseed.com/home.aspx).

### Phenotypic characterization

#### Experimental design

An experiment of 4,608 focal plants of *A. thaliana* and 12,672 neighbor plants was set up at the University of Lille 1 (North, France) in March 2013 using a split-plot design arranged as a randomized complete block design (RCBD) with 12 treatments nested within four blocks. These 12 competition treatments correspond to (Figure 1):

- one control treatment where *A. thaliana* was grown alone (i.e. absence of interaction; hereafter named treatment A).
- 10 interspecific interaction treatments corresponding to the full combination of the three neighboring species *P. annua* (P), *S. media* (S) and *V. arvensis* (V): PPP, SSS, VVV, PPS, PPV, PSS, PVV, SSV, SVV and PSV.
- one intraspecific interaction treatment (hereafter named treatment AAA). This treatment was included in the experiment to test whether the differences observed between the treatment where *A. thaliana* was grown alone (i.e. treatment A) and the treatments of interspecific interactions were not only due to the presence of a neighbor plant, but were rather dependent on either the identity of the neighboring species or the combination of neighboring species. With respect to the barochorous mode of seed dispersal in *A. thaliana* (Weinig et al., 2006; Wender et al., 2005), the intraspecific interaction treatment corresponds to intra-genotypic interaction.

**Figure 1.**
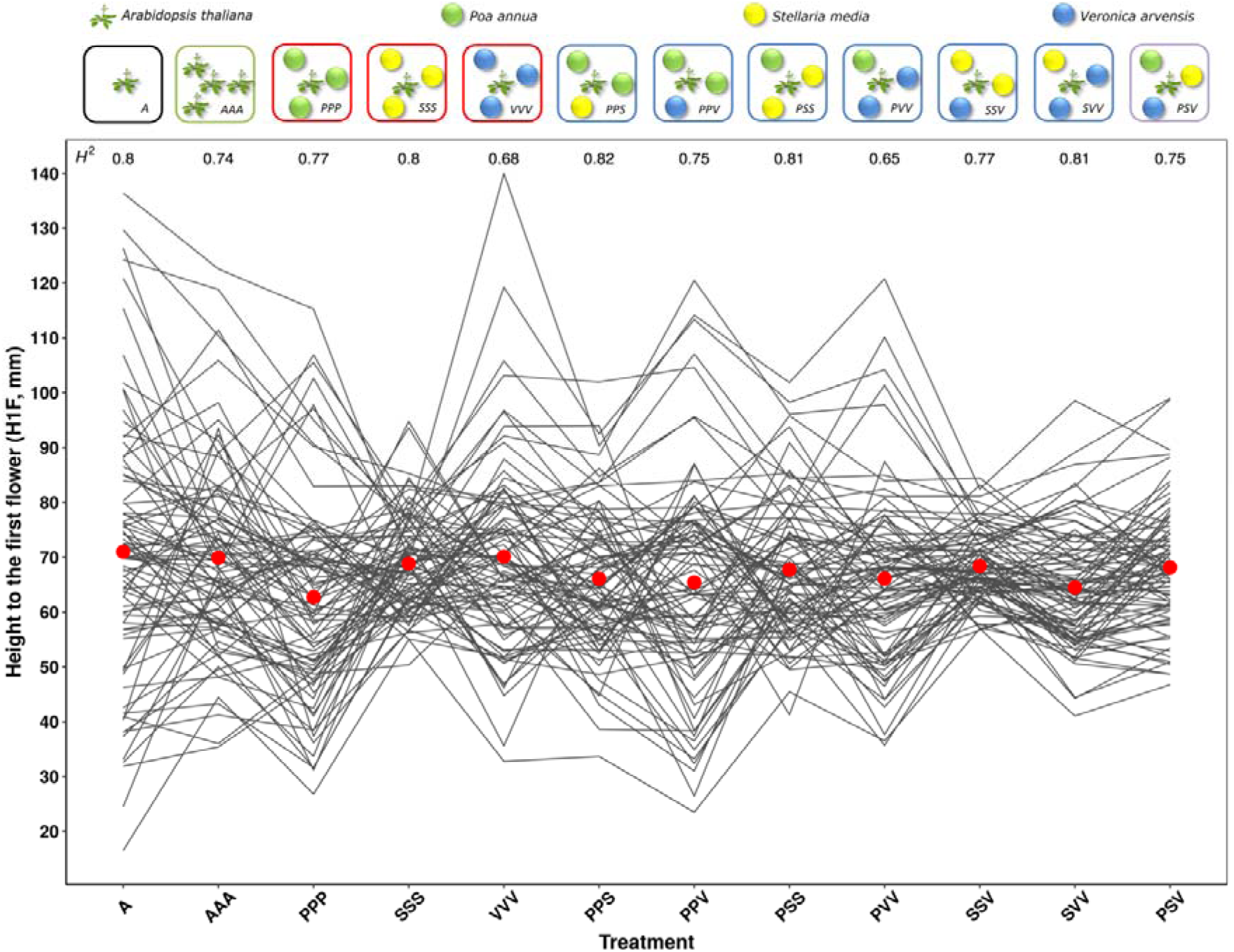
Natural genetic variation of reactions norms of 89 TOU-A accessions across 12 plant-plant interaction treatments. (**a**) Diagram illustrating the 12 treatments. (**b**) Height from the soil to the first flower on *A. thaliana* (H1F). Each line links the genotypic values of one of 89 TOU-A accessions. The two remaining accessions A1-69 and A1-117 are not represented due to missing BLUP values in the PPV and SSS treatments, respectively. For a given treatment, the mean H1F genotypic value among the accessions is represented by a red dot.

Each ‘block x competition treatment’ combination was represented by 96 pots (7 cm x 7 cm x 7 cm, vol. ∼250 cm^3^; TEKU MQC) filled with damp standard culture soil (Huminsubstrat N3, Neuhaus) and disposed in staggered rows, each pot corresponding to one of the 96 TOU-A accessions. The 17^th^ of January 2013 (day 0), a minimum of five *A. thaliana* seeds were sown in the central position of each pot. For all the treatments (with the exception of treatment A), seeds for neighboring plants were evenly spaced, two cm away from the *A. thaliana* central position (Figure 1). Germination date of *A. thaliana* focal seedlings was daily monitored until six days after sowing. At this time, five accessions had a poor germination rate (between 0% and 6.25%) and were therefore discarded from further analyses. Plants that germinated after 6 days (i.e. 1.16%) were also discarded from further analyses.

*A. thaliana* focal seedlings and neighboring seedlings were thinned to one per pot 18 to 20 days after seed sowing. Plants were grown at 20 °C under natural light supplemented by artificial light to provide a 16-hr photoperiod and were top watered without supplemental nutrients. The experience lasted 87 days, from sowing to harvesting of the last plants.

#### Measured phenotypic traits

Three raw phenotypic traits were measured on each focal plant of *A. thaliana* at the time of their flowering, which was measured as the number of days between germination and flowering dates. The first trait corresponds to the height from the soil to the first flower on the main stem (H1F expressed in mm). H1F is related to seed dispersal (Wender et al., 2005) and shade avoidance (Dorn et al., 2000) in *A. thaliana*. The two other traits were used as proxies of resources accumulation. The maximum diameter of the rosette was measured at the nearest millimeter (DIAM; Weinig et al., 2006). This trait is a proxy of the growth of the rosette of the focal plant from germination to flowering. The above-ground dry biomass (BIOMASS, expressed in grams, with a precision down to the tenth of a milligram) was estimated by drying the aboveground portion for 48H at 60°C.

We additionally quantified the strategy adopted by *A. thaliana* in response to neighboring plants by calculating the ratio HD as the height of the first flower on the rosette diameter (i.e. H1F/DIAM). Values of HD below and above 1 would correspond to an aggressive and escape strategy, respectively (Baron et al., 2015).

Plants that had not flowered 87 days after sowing (i.e. 1.15%) were assigned a flowering date value of 87. H1F and HD were therefore not available for these plants.

### Statistical analysis

#### Exploring natural variation of plant-plant interactions at different levels of complexities

The following mixed model (PROC MIXED procedure, REML method, SAS 9.3, SAS Institute Inc) was used to explore the genetic variation of response among the 96 TOU-A accessions:

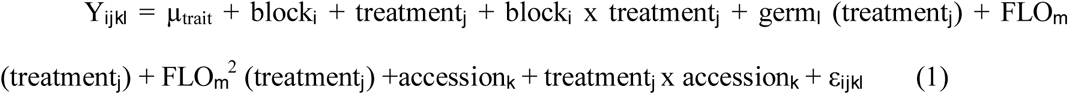

where ‘*Y*’ is one of the phenotypic traits scored on focal *A. thaliana* plants, ‘µ’ is the overall phenotypic mean; ‘block’ accounts for differences in micro-environment among the four experimental blocks; ‘treatment’ corresponds to effect of the 12 treatments (A, AAA, PPP, PPS, PPV, PSS, PSV, PVV, SSS, SSV, SVV and VVV); ‘accession’ measures the effect of the 91 accessions; the interaction term ‘treatment x accession’ accounts for genetic variation in reaction norms across the 12 treatments; the term ‘germ(treatment)’ is a covariate accounting for natural variation for the germination date between the 91 accessions; and ‘ε’ is the residual term. Because phenotypic traits were measured at flowering time, we controlled the putative linear and non-linear effects of this phenological stage by including the linear term ‘FLO’ as well as the quadratic term ‘FLO*FLO’ in model (1).

All factors were treated as fixed effects; with the exception of the term ‘accession’ that was treated as a random effect. For the calculation of *F*-values, terms were tested over their appropriate denominators. Given the split-plot design used in this study, the variance associated with ‘block x treatment’ was used as the error term for testing the ‘block’ and ‘treatment’ effects. Model random terms were tested with likelihood ratio tests of models with and without these effects.

#### Heritability

Based on variance components estimated by REML (PROC VARCOMP procedure in SAS 9.3, SAS Institute Inc.), the broad-sense heritability of each phenotypic trait (*H*^*2*^_trait_) was estimated within each treatment using the following model:

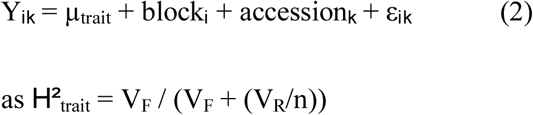

where V_F_ is the estimated between-accession variance component, V_R_ is the residual variance and ‘n’ is the number of replicates per accession.

Significance of *H*^*2*^_trait_ was assessed by testing the significance of the term ‘accession_*k*_’ by fitting model (2) using the PROC MIXED procedure in SAS 9.3 (REML method).

#### Estimation of genotypic values

For each treatment, Best Linear Unbiased Predictors (BLUPs) were obtained for each accession using the following model (PROC MIXED procedure, REML method, SAS 9.3, SAS Institute Inc).

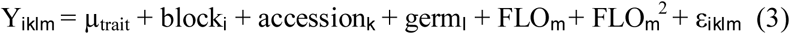

Because *A. thaliana* is a highly selfing species (Platt et al., 2010), BLUPs correspond to genotypic values of accessions.

#### Genetic correlations

For each phenotypic trait, we estimated the strength of ‘accession x treatment’ interactions by estimating across-environment genetic correlations for each pairwise treatment combination. Genetic correlations were estimated by calculating the Pearson correlation coefficient based on BLUPs by using the *cor.test* function implemented in the *R* environment. Significant crossing reaction norms were detected by testing whether 95% confidence intervals of Pearson’s *r* were not overlapping with the value of 1.

To test whether the four phenotypic traits were not redundant, we estimated for each treatment the genetic correlation for each pair of traits, by calculating the Pearson correlation coefficient based on BLUPs as described above.

### Genome wide association mapping

The effects of population structure on phenotype-genotype associations has been demonstrated to be limited in the TOU-A population (Brachi et al., 2013; Baron et al., 2015). Nevertheless, GWA mapping was run using a mixed-model approach implemented in the software EFFICIENT MIXED-MODEL ASSOCIATION EXPEDITED (EMMAX, H.M. Kang et al., 2010). This model includes a genetic kinship matrix as a covariate to control for the effect of the demographic history of the TOU-A population. This kinship matrix was estimated on the whole set of 1,692,194 SNPs detected among the 91 accessions.

In this study, we discarded SNPs with more than 16 missing values across the 91 accessions. In addition, because rare alleles may lead to an inflation of low *p*-values (Atwell et al., 2010; Brachi et al., 2010; H.M. Kang et al., 2010), we only considered SNPs with a minor allele relative frequency (MARF) > 10%, leaving us with 630,234 SNPs.

### Genetic architecture of plant-plant interactions

For each trait, we compared the genetic architecture among the 12 treatments by focusing on the 200 most associated SNPs (i.e. top SNPs) of each treatment. This number of top SNPs represents ∼ 0.03% of the total number of SNPs and has been previously demonstrated to be appropriate and conservative to describe in the TOU-A population the genetic architecture of a set of 29 complex quantitative traits related to phenology, development and fecundity (Frachon et al., 2017). The degree of flexibility of the genetic architecture among the 12 treatments was estimated by calculating the degree of environmental pleiotropy of a given top SNP, which is defined as the number of treatments that shared this top SNP (Wang et al., 2010). Specific comparisons within a subset of treatments were illustrated by Venn diagrams using the *jvenn* online plug-in (Bardou et al., 2014).

### Identification of candidate genes associated with response to monospecific and plurispecific interactions

To identify candidate genes associated with plant-plant interactions, we selected the 20 top SNPs for each ‘trait x treatment’ combination. Following Frachon et al. (2017), we then retrieved all the annotated genes located within or overlapping with a 2kb region around each top SNP, using the TAIR 10 database (https://www.arabidopsis.org/), leaving us with 369 unique candidate genes.

To identify the biological processes involved in response to the different neighborhoods, we first merged the list of candidate genes across the four phenotypic traits according to the four following categories: Control = A; Intraspecific = AAA; Monospecific = PPP, SSS and VVV; Plurispecific = PPS, PSS, PPV, PVV, SSV, SVV and PSV. Based on the MAPMAN classification (Provart & Zhu 2003), the four resulting lists of unique candidate genes were then submitted to the classification superviewer tool (http://bar.utoronto.ca/ntools/cgi-bin/ntools_classification_superviewer.cgi) to identify the biological processes significantly over-represented (*P* < 0.01). Only candidate genes from significantly enriched biological processes were further considered for a functional interpretation of the molecular mechanisms involved in plant-plant interactions in our study.

## Results

### Extent of natural genetic variation of response to monospecific and plurispecific interactions

Highly significant genetic variation was found across the 12 treatments for the four phenotypic traits scored on focal *A. thaliana* plants (Table 1). Similar results were observed without considering the treatments where *A. thaliana* was grown alone or with clones (Tables S1 and S2). Broad-sense heritability values estimated for the four phenotypic traits within each treatment were all highly significant (Supporting information Table S3) and ranged from 0.65 to 0.96 (mean = 0.82, median = 0.81; Supporting information Table S4), suggesting that a large fraction of the phenotypic variation observed within each treatment was driven by genetic differences among the local *A. thaliana* accessions (Figure 1). For each treatment, genetic correlations between the four phenotypic traits were significantly different from the unity (absolute values of Pearson’s *r*: min = −0.72, max = 0.86, mean = −0.02, median = - 0.21), suggesting that the traits scored in this study partly behave independently (Supporting information Figure S1).

**Table 1:** Natural variation of four phenotypic traits scored on *A. thaliana* plants in 12 treatments. Bold *P*-values indicate significant effects after FDR correction. Model random terms were tested with likelihood ratio tests (LRT) of models with and without these effects. Random effects are in italics. H1F: height from the soil to the first flower on the main stem, DIAM: maximum diameter of the rosette, BIOMASS: aboveground dry biomass, HD = H1F / DIAM.

| Model terms | Traits |  |  |  |  |  |  |  |
| --- | --- | --- | --- | --- | --- | --- | --- | --- |
|  | H1F |  | DIAM |  | BIOMASS |  | HD |  |
|  | *F or LRT* | *P* | *F or LRT* | *P* | *F or LRT* | *P* | *F or LRT* | *P* |
| block | 3.51 | **3.27E-02** | 0.54 | 6.65E-01 | 3.69 | **2.86E-02** | 0.70 | 6.01E-01 |
| treatment | 24.37 | **2.55E-32** | 3.34 | **2.10E-04** | 31.35 | **2.55E-32** | 1.46 | 1.64E-01 |
| *accession* | 969.70 | **9.70E-212** | 821.20 | **1.24E-179** | 556.70 | **3.07E-122** | 1094.40 | **1.53E-238** |
| *treatment\*accession* | 14.10 | **2.55E-04** | 39.40 | **6.04E-10** | 174.90 | **3.53E-39** | 33.20 | **1.37E-08** |
| germ(treatment) | 1.61 | 1.01E-01 | 0.79 | 6.65E-01 | 2.42 | **5.59E-03** | 0.90 | 6.01E-01 |
| flo(treatment) | 53.12 | **2.55E-32** | 9.56 | **4.66E-18** | 52.68 | **2.55E-32** | 8.70 | **4.02E-16** |
| flo*flo(treatment) | 39.04 | **2.55E-32** | 7.59 | **1.06E-13** | 18.26 | **2.55E-32** | 6.97 | **2.54E-12** |

Importantly, as evidenced by highly significant ‘Treatment x Accession’ interactions, strong genetic variation of reaction norms was found for the four phenotypic traits, with or without considering the treatments where *A. thaliana* was grown alone or with clones (Table 1, Supporting information Tables S1 and S2). Across the four phenotypic traits, across-environment genetic correlations ranged from 0.19 to 0.86 (mean = 0.57, median = 0.57; Figure 2), indicating that the rank of accessions largely differed among the 12 treatments. This pattern of crossing reaction norms is well illustrated for the height from the soil to the first flower (Figure 1).

**Figure 2.**
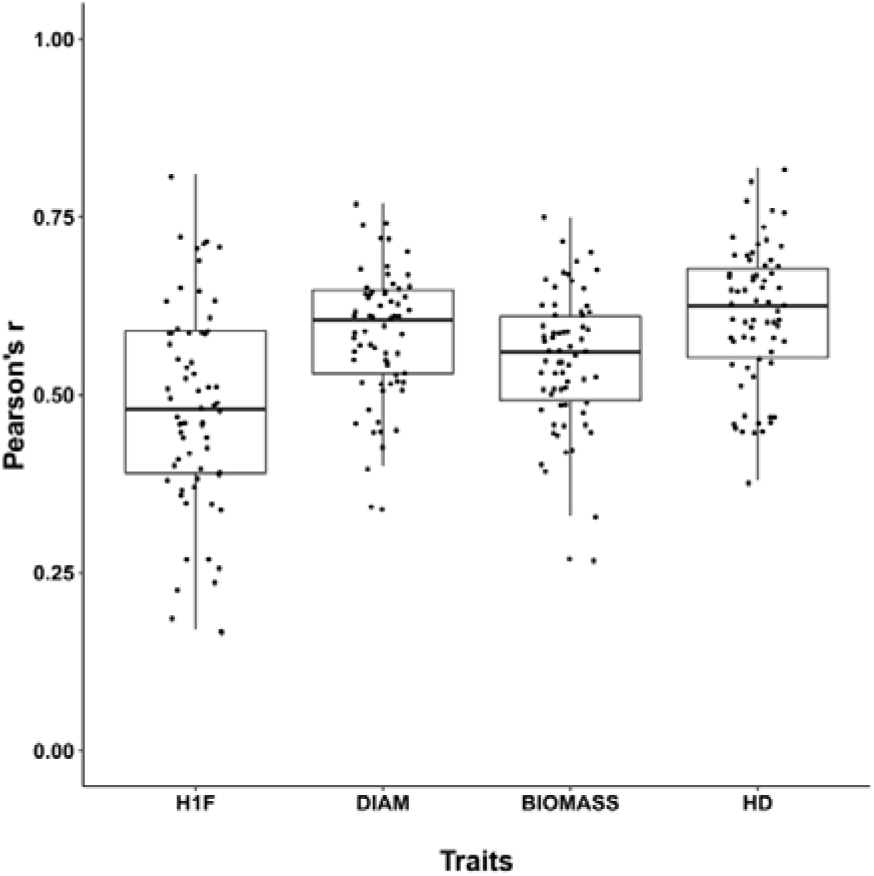
Pairwise genetic correlation coefficients of Pearson among the 12 treatments for each phenotypic trait. The dots correspond to the 66 pairwise treatment combinations.

### Genetic architecture revealed by GWA mapping

To compare the genetic architecture underlying the competitive response of *A. thaliana* between monospecific and plurispecific neighborhoods, we used GWA mapping based on 630,234 SNPs (i.e. 1 SNP every 189bp). Following the methodology of a previous GWAS performed on the local TOU-A population (Frachon et al., 2017), we described the genetic architecture by extracting for each treatment the 200 top SNPs associated with each of the four phenotypic traits (Supporting information Figure S2), leading to a final set of 4091 unique SNPs.

Across the 12 treatments, the degree of environmental pleiotropy of a given top SNP followed an L-shaped distribution (Figure 3). For the traits H1F, DIAM and HD, more than 75.6% of top SNPs were specific to a single treatment, indicating that the genetic bases are largely distinct among the 12 treatments (Figure 3). The genetic architecture was less flexible among the 12 treatments for the trait BIOMASS with (i) the detection of less than 52% of top SNPs being specific to a single treatment and (ii) the identification of top SNPs shared among up to eight treatments (Figure 3). This latter observation likely resulted from the detection of a common association peak at the beginning of chromosome 4 between the 12 treatments (Supporting information Figure S2).

**Figure 3.**
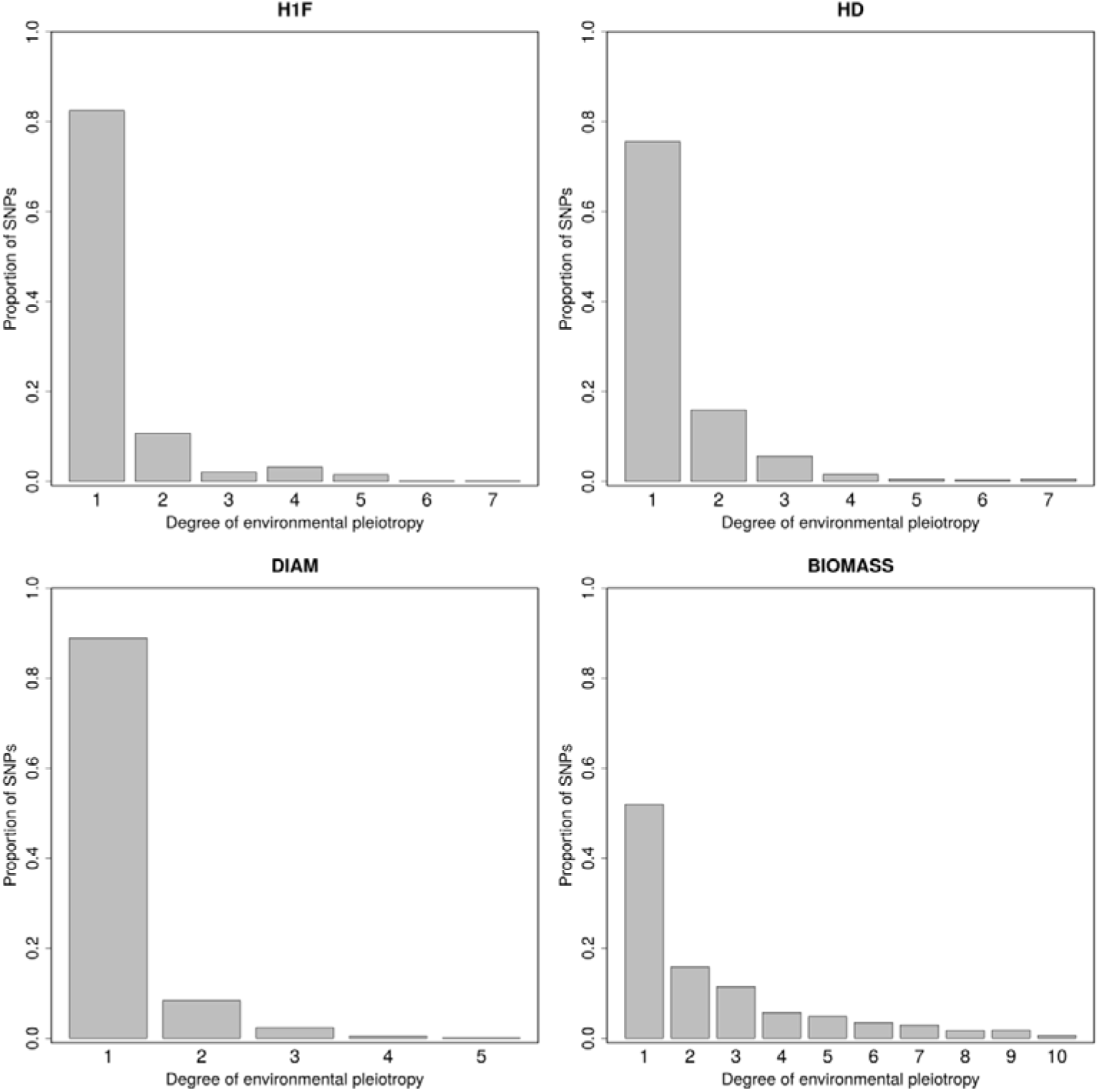
Degree of flexible genetic architecture of *A. thaliana* among the 12 treatments when considering a threshold of 200 top SNPs. For each phenotypic trait, bar plots represent the frequency distribution of the number of treatments that share a top SNP. H1F: height from the soil to the first flower on the main stem, DIAM: maximum diameter of the rosette, BIOMASS: aboveground dry biomass, HD = H1F / DIAM.

Therefore, in this study, the genetic architecture largely depends on both the composition and assemblage of the neighborhood of *A. thaliana*. Firstly, for the traits H1F, DIAM and HD, the genetic architecture of competitive response of *A. thaliana* to monospecific interactions was highly dependent on the identity of the neighboring species (Supporting information Figures S3, S4 and S5). As illustrated for HD, on average 79% of top SNPs were specific to either the control treatment or one of the three monospecific interaction treatments (Figure 4). On the other hand, less than half of top SNPs (i.e. 44.5%) associated with BIOMASS were specific either the control treatment or one of the three monospecific interaction treatments (Supporting information Figure S6).

**Figure 4.**
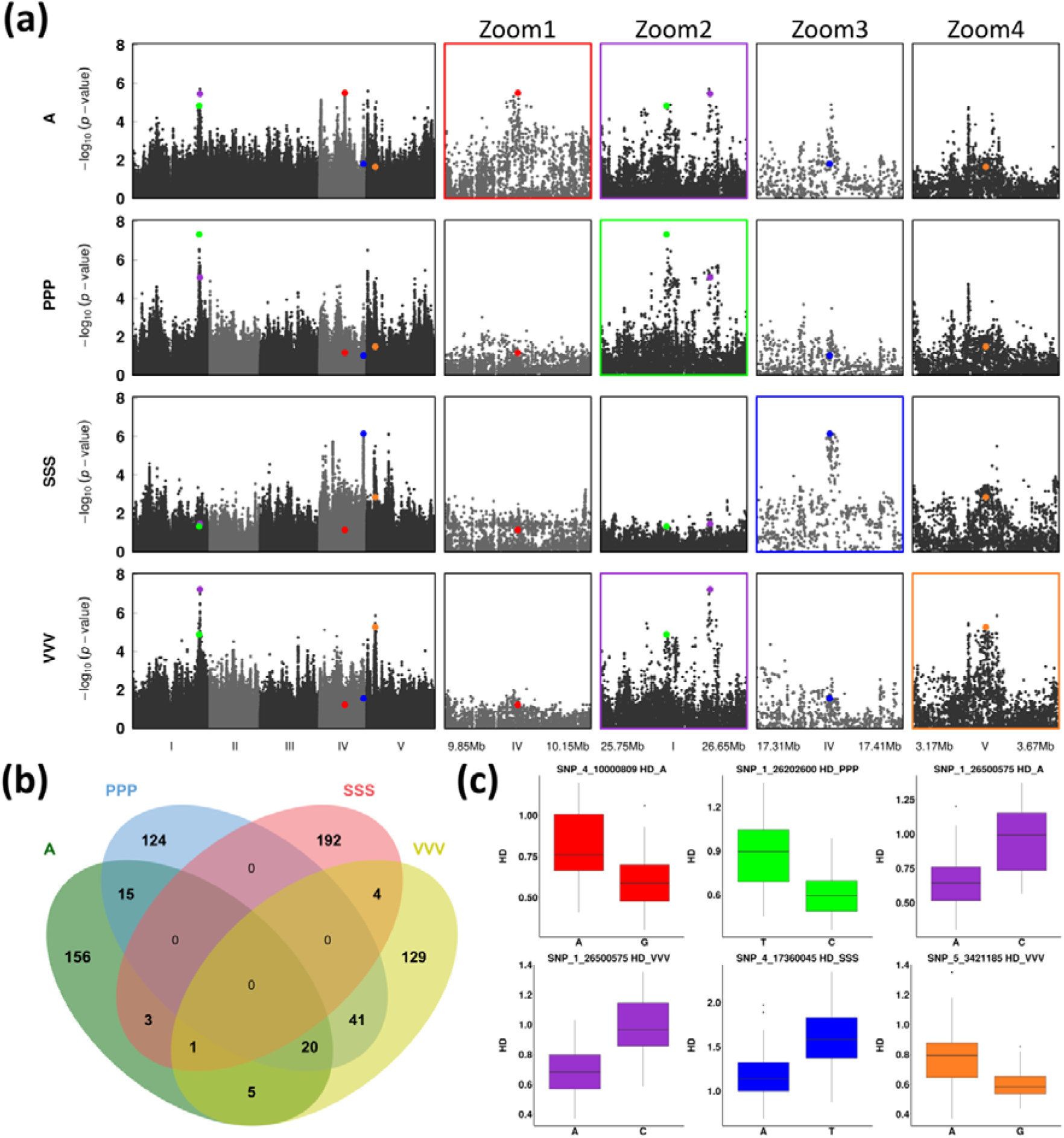
Identification of genomic regions associated with monospecific interactions for the ratio ‘height of the first flower / the rosette diameter’ (HD) in the TOU-A population. (**a**) Left panel: Manhattan plots of GWA mapping results for the A, PPP, SSS and VVV treatments. The *x*-axis indicates the physical position of the 630,234 SNPs along the five chromosomes. The *y*-axis indicates the −log_10_ *p*-values using the mixed model implemented in the software EMMAX using SNPs with MARF > 10% and missing data < 75%. Mid-panel and right panel: zooms on top SNPs illustrating the degree of specificity of genetic architecture among the treatments A (zoom1, red dot; zoom2, purple dot), PPP (zoom2, green dot), SSS (zoom3, blue dot) and VVV (zoom4, orange dot). (**b**) Venn diagram partitioning the HD SNPs detected among the lists of 200 top SNPs for the A, PPP, SSS and VVV treatments. **(c)** Box-plots illustrating the effects of the five top SNPs colored in panel (a) in their respective treatment.

Secondly, the genetic architecture of competitive response of *A. thaliana* was also highly dependent on the three-way combination of *P. annua, S. media* and *V. arvensis*. In particular, for the traits H1F, DIAM and HD, more than 93% of top SNPs associated with the response of *A. thaliana* to the simultaneous presence of *P. annua, S. media* and *V. arvensis* were not present in the sets of 200 top SNPs identified in the three monospecific interaction treatments (Supporting information Figures S3, S4 and S5). For example, a very neat peak of association for HD was identified at the beginning of chromosome 3 in the treatment PSV but not in the treatments PPP, SSS and VVV (Figure 5a).

**Figure 5.**
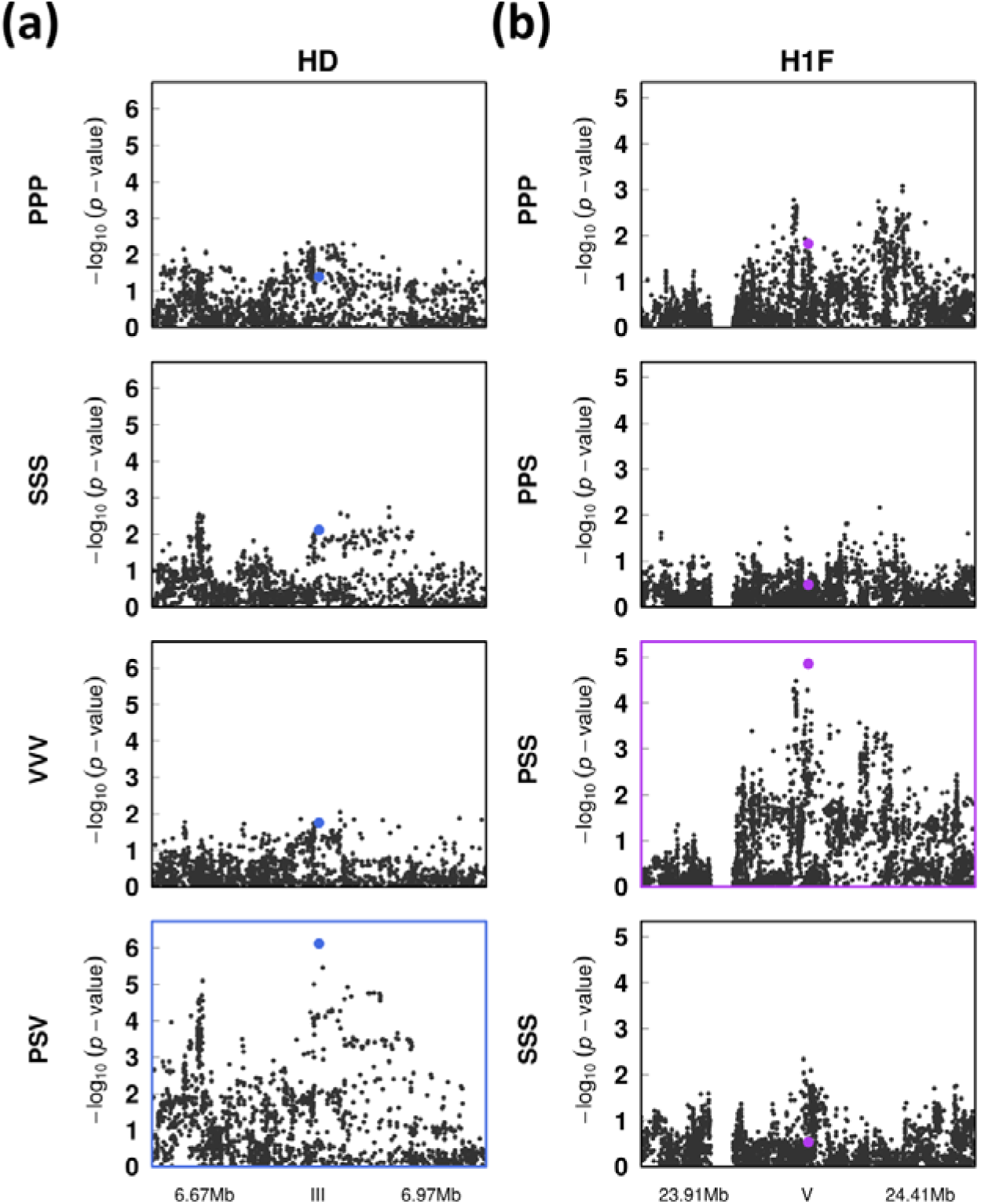
Comparison of the genetic architecture between monospecific and plurispecific interactions in the TOU-A population. (**a**) Zooms on an association peak identified for the ratio ‘height of the first flower / the rosette diameter’ (HD), which is specific to the plurispecific interaction treatment PSV. (**b**) Zooms illustrating an association peak identified for the height from the soil to the first flower (H1F), which depends on the assemblage between the neighboring species *P. annua* and *S. media*. The *x*-axis indicates the physical position of the SNPs along the considered genomic region. The *y*-axis indicates the −log_10_ *p*-values using the mixed model implemented in the software EMMAX using SNPs with MARF > 10% and missing data < 75%.

Thirdly, the identity of top SNPs associated with the response of *A. thaliana* to the presence of a specific pair of neighboring species largely differed not only from the top SNPs identified in the corresponding monospecific interaction treatments (as previously observed for the treatment PSV), but also between the two assemblages based on this pair of neighboring species (Supporting information Figure S2). In the latter case, less than 16% of top SNPs were shared between treatments with the same composition (i.e. two neighboring species) but with different assemblages (i.e. two plants of species A + 1 plant of species B *vs* one plant of species A + two plants of species B) (Supporting information Figure S2). As an illustration, we detected a neat association peak for H1F at the end of chromosome 5 in the treatment PSS (i.e. one *P. annua* individual + two *S. media* individuals) but neither in the treatment PPS (i.e. two *P. annua* individuals + one *S. media* individual), nor in the treatments PPP and SSS (Figure 5b).

### Identification of enriched biological processes and underlying candidate genes

We retrieved 369 unique genes located within or overlapping a 2kb region around the 20 top SNPs of each ‘trait x treatment’ (Dataset1). Considering this entire set of genes, only the ‘transport’ class was significantly over-represented in frequency compared to the overall class frequency in the *Arabidopsis thaliana* MapMan annotation (Normed Frequency = 1.87, Supporting information Table S5), whereas the ‘DNA’ and ‘not assigned’ classes were significantly under-represented (Normed Frequency = 0.17 and 0.86, respectively, Supporting information Table S5).

Among the four interaction categories (i.e. Control, Intraspecific, Monospecific and Plurispecific), we observed strong differences in the number and identity of enriched biological processes (Supporting information Table S6). No biological process was found significantly enriched for the ‘Control’ and ‘Intraspecific’ interaction categories (Supporting information Table S6). In contrast, significantly enriched biological processes were detected in the context of interspecific interactions. For the ‘Monospecific’ interaction category, we detected a significant enrichment for the ‘tetrapyrrole synthesis’ class, which was represented by the genes *HEME OXYGENASE 3* (*HO3*) and *HEMB1* (Supporting information Table S7), both identified for HD in the PPP treatment. For the ‘Plurispecific’ interaction category, we identified three significantly enriched biological processes, i.e. signaling, transport and DNA with 18, 16 and 5 underlying candidate genes, respectively (Supporting information Table S6). In the ‘signaling’ class, 12 out of the 18 candidate genes encode receptor-like kinases (RLKs), i.e. one wall-associated receptor kinase (WAK), two proline-rich extensin-like receptor kinases (PERKs), two cystein-rich receptor-like kinases (CRKs) and seven leucine-rich repeat receptor-like protein kinases (LRR-RLKs) (Supporting information Table S6). The six remaining proteins were related to signaling (EPS15 HOMOLOGY DOMAIN 1 (EHD1), NO POLLEN GERMINATION 1 (NPG1) and IQ-DOMAIN 17 (IQD17)), a G-box family protein (G-BOX REGULATING FACTOR 6, GRF6), an exordium like protein (EXORDIUM LIKE 2, EXL2) and the RPM1-interacting protein 4 family protein (AT5G40645). In the ‘transport’ class, we identified several genes encoding diverse transporters (i) four ATP-BINDING CASSETTE transport proteins (ABC transporters), which were P-GLYCOPROTEIN 18 (PGP18), ATP-BINDING CASSETTE A3 (ABCA3), ATP-BINDING CASSETTE F2 (ABCF2) and PLEIOTROPIC DRUG RESISTANCE 13 (PDR13) and (ii) two homologous phosphate transport proteins (PHOSPHATE 1 (PHO1) and its homolog PHO1;H1). We also identified four proteins related to the transport of mineral nutrients, i.e. one protein related to copper transport (COPPER TRANSPORTER 3, COPT3), two proteins related to magnesium transport (MAGNESIUM TRANSPORTER 4 (MRS2-3) and a magnesium transporter CorA-like family protein (*AT5G09710*)) and a nitrate transporter (NRT1 PTR FAMILY 5.13, NPF5.13). The six other genes are related to the transport of purine, calcium, sugars and peptides (Supporting information Table S7). In the ‘DNA’ class, the five candidate genes correspond to the DNA topoisomerase VI sub-unit A SPORULATION 11-1 (SPO11-1), a DNA glycosylase (AT3G50880), the RNA HELICASE-LIKE 8 (RH8), the DNA LIGASE IV (LIG4) and a histone superfamilly protein (AT5G59970).

## Discussion

Because a focal plant rarely interacts with only one neighboring species either in crop fields or in more natural environments, the genetics of plant-plant interactions need to be studied in a community context. In this study, we compared the genetic architecture of the competitive response of *A. thaliana* between monospecific and plurispecific neighborhoods. To achieve this goal, we adopted a GWA mapping approach combined with the modern standards of ecological genomics. Indeed, the geographical scale at which selective agents act on a species should determine the mapping population used to identify genomic regions associated with ecologically relevant trait phenotypic variation (Bergelson & Roux 2010; Brachi et al., 2013; Roux & Bergelson 2016). Because plants interact with neighbors over short distances, we focused on a highly genetically polymorphic local population of *A. thaliana* known to interact *in situ* with the three neighboring species considered in this study.

### A flexible genetic architecture for competitive response between monospecific and plurispecific neighborhoods

Several studies reported extensive genetic variation of the competitive ability of *A. thaliana* in the context of pairwise heterospecific interactions, both at the worldwide and local scales (Bossdorf et al., 2009; Brachi et al., 2012; Bartheleimer et al., 2015; Baron et al., 2015; Frachon et al., 2017). Based on four phenotypic traits related to resource accumulation and life-history trait such as seed dispersal (Reboud et al., 2004; Wender et al., 2005), we also found extensive local genetic variation of the competitive ability of *A. thaliana* in all the plurispecific neighborhoods tested in this study. More importantly, we detected strong crossing reaction norms not only among the three monospecific interaction treatments, but also among the different plant assemblages surrounding the focal *A. thaliana* accessions. Altogether, these results suggest that the simultaneous interactions of *A. thaliana* with several plant partners can promote maintenance of the high genetic diversity observed in the TOU-A local population (i.e. only 5.6 times less than observed in a panel of 1,135 worldwide accessions) (Frachon et al., 2017). This diversity may in turn confer a high potential for *A. thaliana* to respond to future modifications of the assemblages in the TOU-A plant community.

In agreement with the strong crossing reaction norms observed among the three monospecific interaction treatments, a GWA mapping approach reveals that the genetic architecture of *A. thaliana* competitive response in monospecific neighborhoods was highly dependent on the identity of the neighbor species. Similar results were observed (i) in *A. thaliana* challenged with four neighboring species (including the three species used in this study) in field conditions (Baron et al., 2015) and (ii) in *Oryza sativa* challenged with the three weed species *Echinochloa oryzicola, Monochoria korsakowii* and *Schoenoplectus juncoides* in greenhouse conditions (Onishi et al., 2018). Altogether, these results suggest that the effect of the identity of the neighboring species on the genetic architecture of competitive response is conserved among focal plant species and across diverse phenotyping environments. The genetic architecture of *A. thaliana* competitive response was also highly flexible between monospecific and plurispecific neighborhoods, suggesting that the genetic response of a particular accession of *A. thaliana* in a plurispecific neighborhood can be hardly predicted from the additivity of its genetic responses observed in the corresponding monospecific neighborhoods.

Three non-exclusive and interconnected hypotheses can be proposed to explain the emergence of new QTLs in plurispecific neighborhoods (Figure 6). They are based on (i) the putative generation of new signals or elimination/modification of pre-existing signals (e.g. light, aerial volatile organic compounds, root exudates and nutrient availability) in the context of plurispecific interactions, and (ii) their perception by *A. thaliana*. Firstly, the amount of some signals produced by the neighboring species A is strongly reduced in a plurispecific neighborhood, thereby leading to the strong reduction of the perception/signaling events activated in the context of monospecific interactions. Secondly, the production of new or modified signals in a given neighboring species is triggered by the presence of another neighboring species. Thirdly, the simultaneous presence of different signals produced by the neighboring species A and B leads to the generation of a new signal. The two latter cases correspond to the production of new signals that emerged from higher-order interactions among the neighboring species (Levine et al., 2017). Natural variation in genes involved in the perception, signaling and genetic program triggered by a new set of signals can explain the emergence of new QTLs in plurispecific neighborhoods (Figure 6).

**Figure 6.**
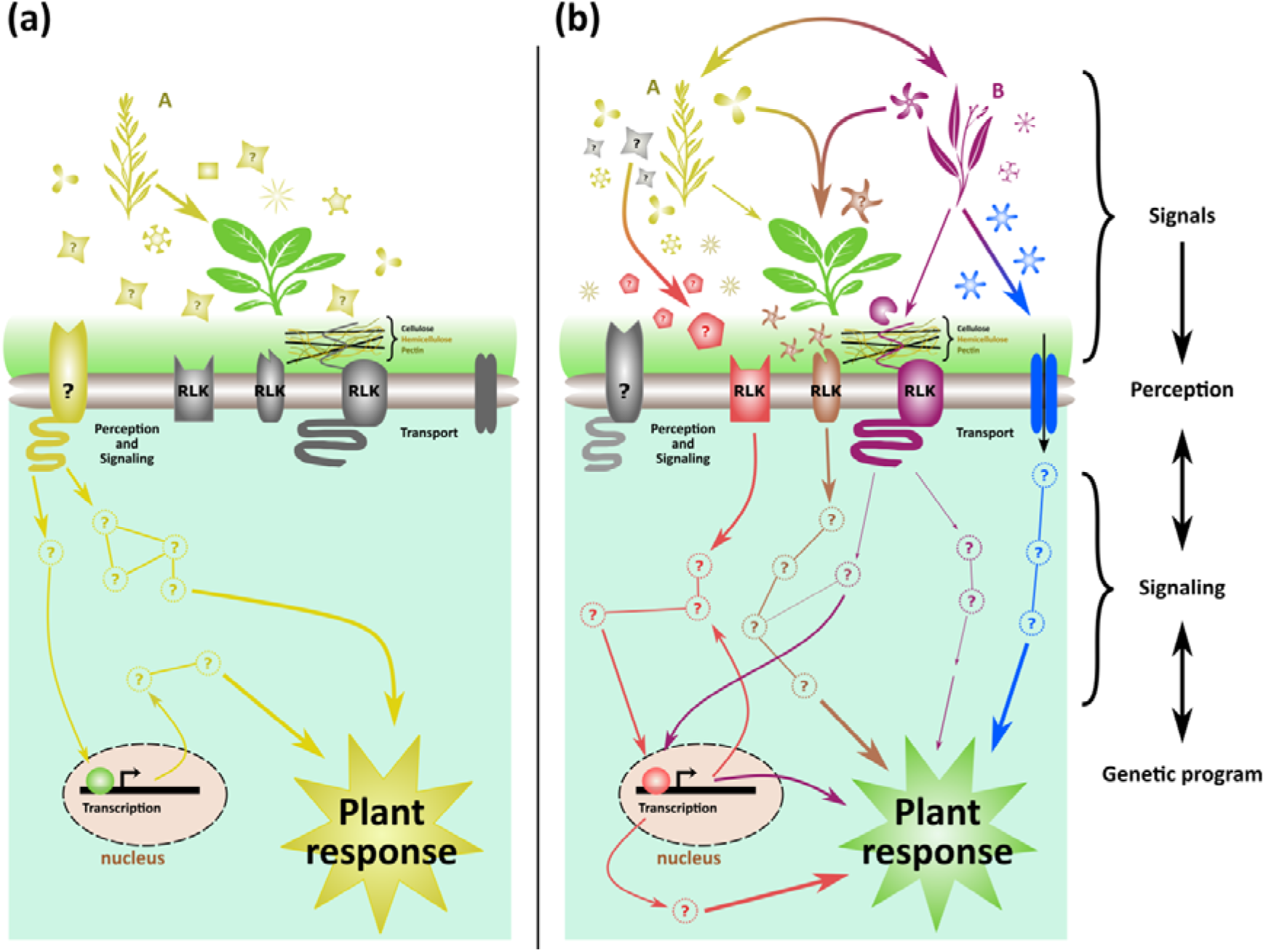
A schematic model to explain the differential identification of QTLs between mono- and plurispecific interactions. **(a) Identification of QTLs in a monospecific neighborhood.** In this diagram, the species A (yellow) produces different compounds (yellow symbols). For example, one of them is perceived by a cell wall receptor of the focal plant (green). The perception of this signal triggers the activation of different signaling pathways. These putative pathways lead to a specific plant response that could be variable among accessions due to genetic diversity of mechanisms underlying perception and/or signaling and/or gene expression events. **(b) Identification of different QTLs in a plurispecific neighborhood.** To explain the identification of different QTLs in response to a plurispecific neighborhood, we propose non-exclusive scenarios that are based on the putative generation of new signals or elimination/modification of pre-existing signals. (i) The amount of some signals produced by the species A could be strongly reduced in presence of species B (purple) leading to a strong reduction of the perception/signaling events occurring through the receptor (yellow) identified in (a). (ii) Another possibility could be that the species A, in response to species B, modifies a signal(s) present in the context of monospecific interactions into a new signal (red), which is perceived and transduced by a receptor like kinase (RLK) leading to a specific plant response. (iii) Likewise, the species B might produce a new signal (signal not produced in the context of monospecific interactions, in blue) that activates a transmembrane transporter leading to a specific plant response. (iv) A fourth case could be the simultaneous presence of different signals produced by the neighboring species A and B that together create a new signal (in brown), which is also perceived and transduced leading to a specific plant response. All these examples require (i) generation and /or modification and /or elimination of signals, and (ii) genetic variation in at least one of the three following mechanisms: perception, signaling and genetic programming. Vector plant patterns have been retrieved from the Vecteezy.com website.

### Biological pathways and candidate genes associated with competitive response depend on the number of neighboring species

While significant enriched biological processes were identified in the context of interspecific interactions, no biological process was found significantly over-represented for the ‘Control’ and ‘Intraspecific’ categories. Technically, this observation might be explained by the fact that multiple functions are involved in such interactions, and/or most of the implicated genes correspond to unknown functions (26/92, Supporting information Table S6), leading to the absence of enrichment of any biological process.

Only the tetrapyrrole biosynthesis pathway represented by the genes *HO3* and *HEMB1* was found for the ‘Monospecific’ interaction category. In plants, tetrapyrroles play essential roles in photosynthesis, respiration, and signal transduction (Mochizuki *et al.*, 2010; Tanaka *et al.*, 2011). The tetrapyrrole biosynthesis pathway consists in two main branches, i.e. the chlorophyll and heme branches. In the chlorophyll biosynthetic pathway, two homologous transcription factors essential for phyA signaling, FAR-RED ELONGATED HYPOCOTYL 3 (FHY3) and FAR-RED IMPAIRED RESPONSE 1 (FAR1), activate *HEMB1* that in turn regulates chlorophyll biosynthesis and seedling growth (Tang *et al.*, 2012). *HO3* encodes a haem oxygenase protein. Haem oxygenases have recently emerged as players in plant cell protection to oxidative damage. In this context, *HO3* is involved in salinity tolerance in *A. thaliana* by controlling K+ retention (Bose *et al.*, 2013). Altogether, these findings suggest that photosynthesis, and more widely light perception and signaling, might be essential in the case of monospecific interactions.

In the ‘Plurispecific’ interaction category, the major over-represented biological process was related to signaling processes and mainly composed by RLKs (receptor like kinases). Because RLKs can bind a large variety of ligands, they play essential roles in many plant processes such as plant immunity, development and growth (Tang *et al*., 2017). RLKs tend to be significantly over-represented among genes up-regulated under abiotic (UV-B, wounding and osmotic stress) and biotic (symbiotic or pathogenic interactions) stress conditions (Lehti-Shiu et al., 2009; Tang *et al*., 2017). Our findings suggest that RLKs might also be key players in plant-plant interactions. Several of our candidate RLKs have been reported to be essential in plant development and morphogenesis. TMK1 (TRANSMEMBRANE KINASE 1) interacts with the auxin binding protein ABP1 and activates plasma membrane–associated ROPs (Rho-like guanosine triphosphatases (GTPase)), which control cytoskeleton modifications and the shape of leaf pavement cells in *A. thaliana* (Dai et al., 2013; Xu et al., 2014). IKU2 (HAIKU2) controls endosperm proliferation and seed size (X. Kang et al., 2013). While RUL1 (REDUCED IN LATERAL GROWTH 1) is involved in secondary root growth (Agusti et al., 2011), PERK13 (PROLINE EXTENSIN LIKE RLK) acts as a negative regulator of root hair growth (Y. Hwang et al., 2016). Some other candidate RLKs detected in our study are related to abiotic stress responses: RPK1 is involved in a protein complex governing superoxide production and signaling at the cell surface and controlling senescence and cell death (Koo *et al*., 2017); CRK8 is transcriptionally regulated in response to light stress and ozone (Wrzaczek et al., 2010) and EHD1 has been shown to confer salt tolerance when it is over-expressed in *A. thaliana* (Bar *et al.*, 2013). All these functions and others still unknown might participate to plant response to competition either *via* developmental or stress responses. Identification of the corresponding ligands together with mutant phenotyping in the context of plant-plant interactions should shed some light on the molecular mechanisms underlying these interactions.

The second major over-represented biological process in the ‘Plurispecific’ interaction category is related to transport functions. Interestingly, 4 out of the 16 transport related proteins correspond to ABC transporter proteins. Although none of them have been functionally characterized, these transporters have been recently proposed as key players of plant adaptation to their abiotic or biotic environment (J.U. Hwang *et al*., 2016). Because of their diverse substrate specificities, they might constitute essential components of perception/signaling pathways activated during plurispecific plant-plant interactions Moreover, two proteins related to phosphate transport, PHO1 and its homolog PHO1;H1, were identified. The *pho1* mutant of *A. thaliana* exhibits inorganic phosphate (Pi) deficiency in the Pi export from roots to shoots, resulting in strong Pi deficiency in above-ground tissues (Hamburger et al., 2002). *PHO1;H1* can complement the *pho1* mutant revealing some functional redundancy between these two proteins (Stefanovic et al., 2007). Interestingly, *PHO1* has been identified in a genome-wide association study as a candidate gene underlying natural variation in root architecture and shown to be involved in lateral root plasticity response *via* its interplay with different signals (Rosas *et al*., 2013). This finding is particularly interesting in regard to the potential role of root architecture and Pi signaling in plant development in the context of interspecific interactions. Finally, the identification of candidate genes associated with copper (*COPT3*), magnesium (*MRS2-3* and *AT5G09710*) and nitrate (*NPF5.13*, Léran et al., 2014) transport indicates that nutrient foraging might also be a major response strategy in the context of plurispecific plant-plant interactions (Pierik et al., 2013).

Subrahmaniam *et al.* (2018) reported seven categories of functions previously identified in artificial environments simulating plant–plant interactions: photosynthesis, hormones, cell wall modification and degradation, defense against pathogens, ABC (ATP-binding cassette) transporters, histone modification and meristem identity/life history traits. Surprisingly, only 39 genes (out of our 369 candidate genes) are related to these functional categories (Supporting information Table S8), highlighting the added value of challenging focal plants directly with neighbor plants to identify the molecular mechanisms underlying neighbor perception, signaling and the resulting cell reprogramming.

In conclusion, our results suggest that plants can integrate and respond to different species assemblages depending on the identity and number of each neighboring species, through a large range of genes associated mainly with perception and signaling processes leading to developmental and stress responses (Figure 6). Complementarily to our GWA study, transcriptomic and proteomic analyses of *A. thaliana* plants exposed to monospecific and plurispecific neighborhoods would help to identify genes and proteins that are differentially regulated under these conditions. To our knowledge, no such studies comparing global changes in protein and gene expression between monospecific and plurispecific neighborhoods have been reported so far (Subrahmaniam et al., 2018). Another step to understand the mechanisms underlying natural variation of plant-plant competitive responses would be (i) to functionally validate the identified candidate genes. This would open the way to functional analyses to investigate (ii) the nature of the signals perceived by the plant, and (iii) decipher the signaling pathways resulting from signal perception, leading to the plant response.

## Acknowledgments

Special thanks are given to Cédric Glorieux, Nathalie Faure and Angélique Bourceaux for their assistance during the common garden experiment. This work was funded by a PhD fellowship from the University of Paul Sabatier Toulouse and a PhD fellowship from the University of Lille 1 – Région Nord-Pas-de-Calais to EB. This study was also supported by the LABEX TULIP (ANR-10-LABX-41; ANR-11-IDEX-0002-02).

## Author contributions

F.R., L.A. and D.R. supervised the project. E.B., L.A. and F.R. designed the experiments. E.B. and J.L. conducted the greenhouse experiment. J.L. and E.B measured the phenotypic traits. C.L. analyzed the phenotypic traits. C.L. performed the GWA mapping. C.L. performed and analyzed the enrichment tests. C.L., D.R. and F.R. wrote the manuscript. All authors contributed to the revisions.

