## Supporting information for "The genomic architecture of competitive response of *Arabidopsis thaliana* is highly flexible between monospecific and plurispecific neighborhoods"

1 Supplementary Dataset

8 Supplementary Tables

6 Supplementary Figures

**Figure S1. Pairwise genetic correlation coefficients of Pearson among the four phenotypic traits for each treatment.** The dots for each pair of phenotypic traits correspond to the 12 treatments.

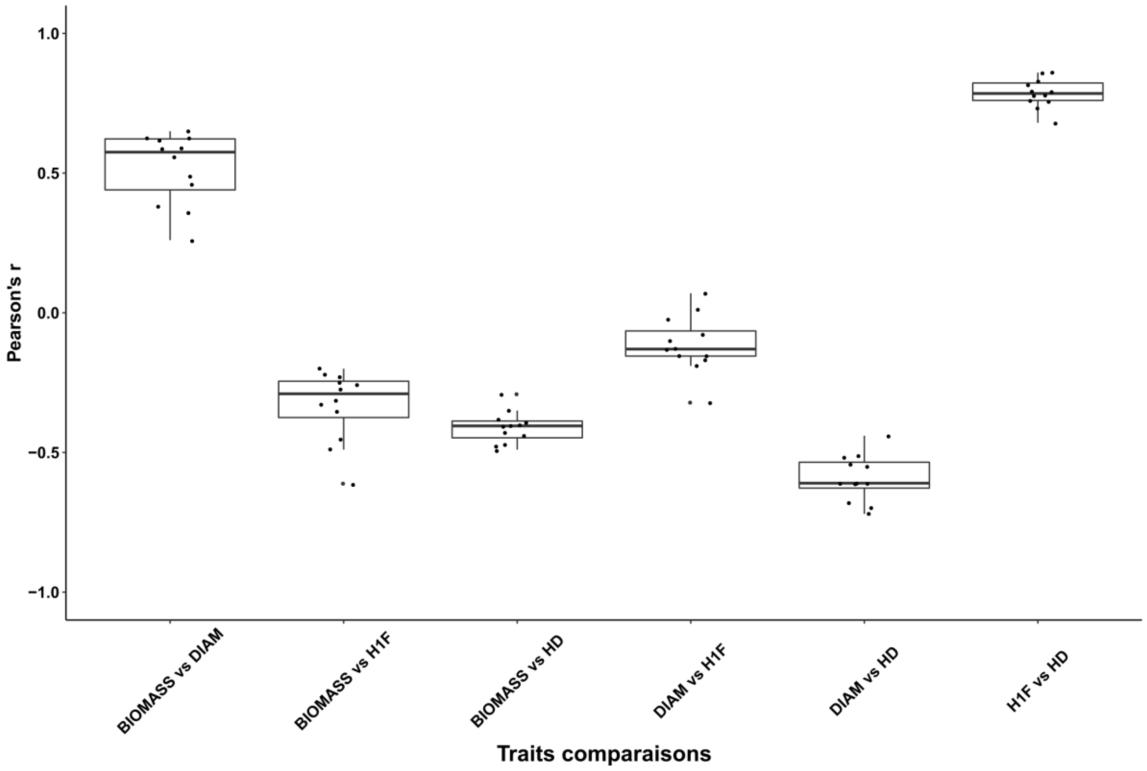

**Figure S2. Identification of genomic regions associated with the four phenotypic traits scored on *A. thaliana* plants in 12 treatments.** The *x*-axis indicates the physical position along the chromosome. The *y*-axis indicates the -log$^{10}$ *p*-values using the EMMAX method. MARF > 10%. On each Manhattan plot, the 200 top SNPs are highlighted in red.

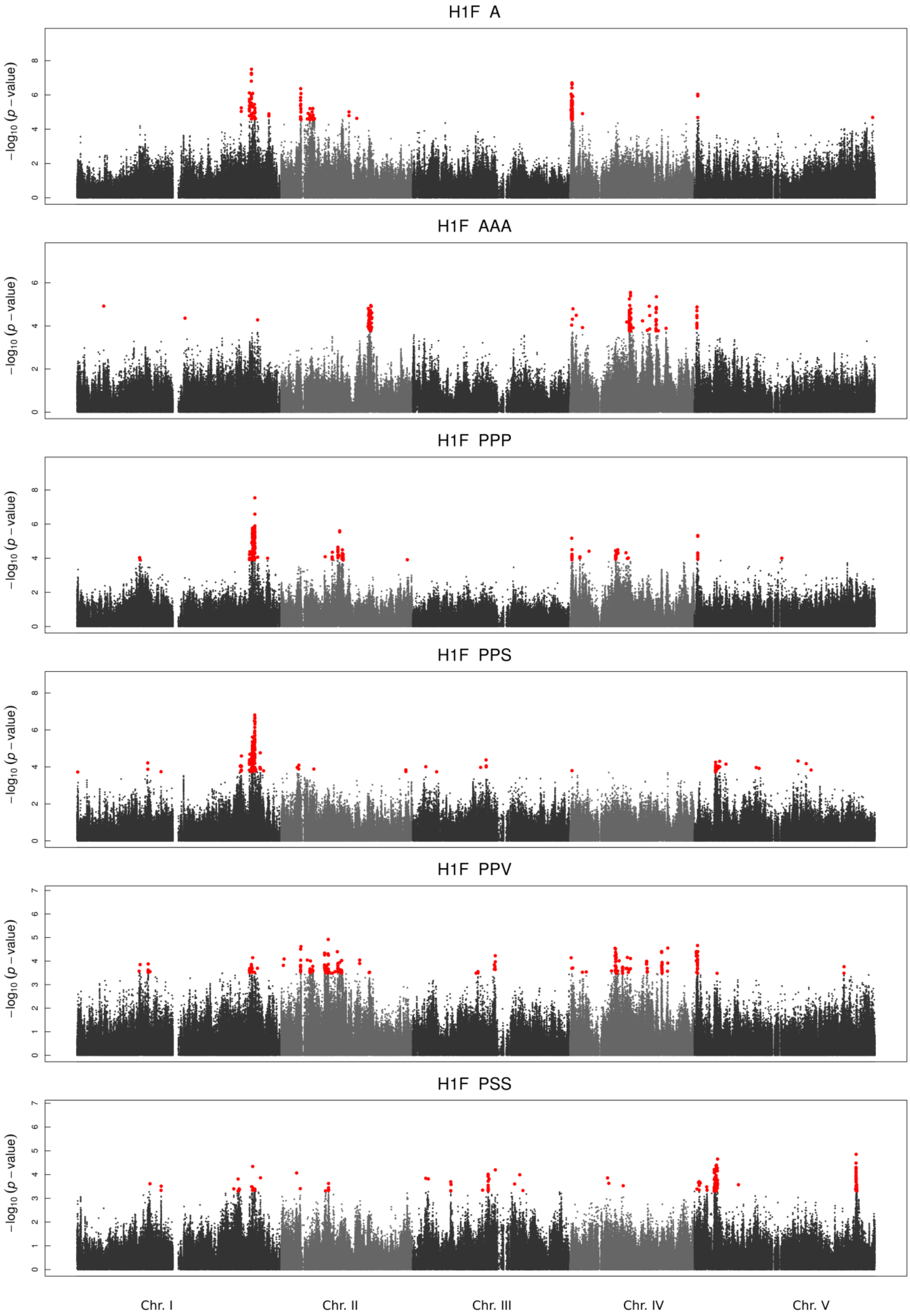

**Figure S2 (continued)**

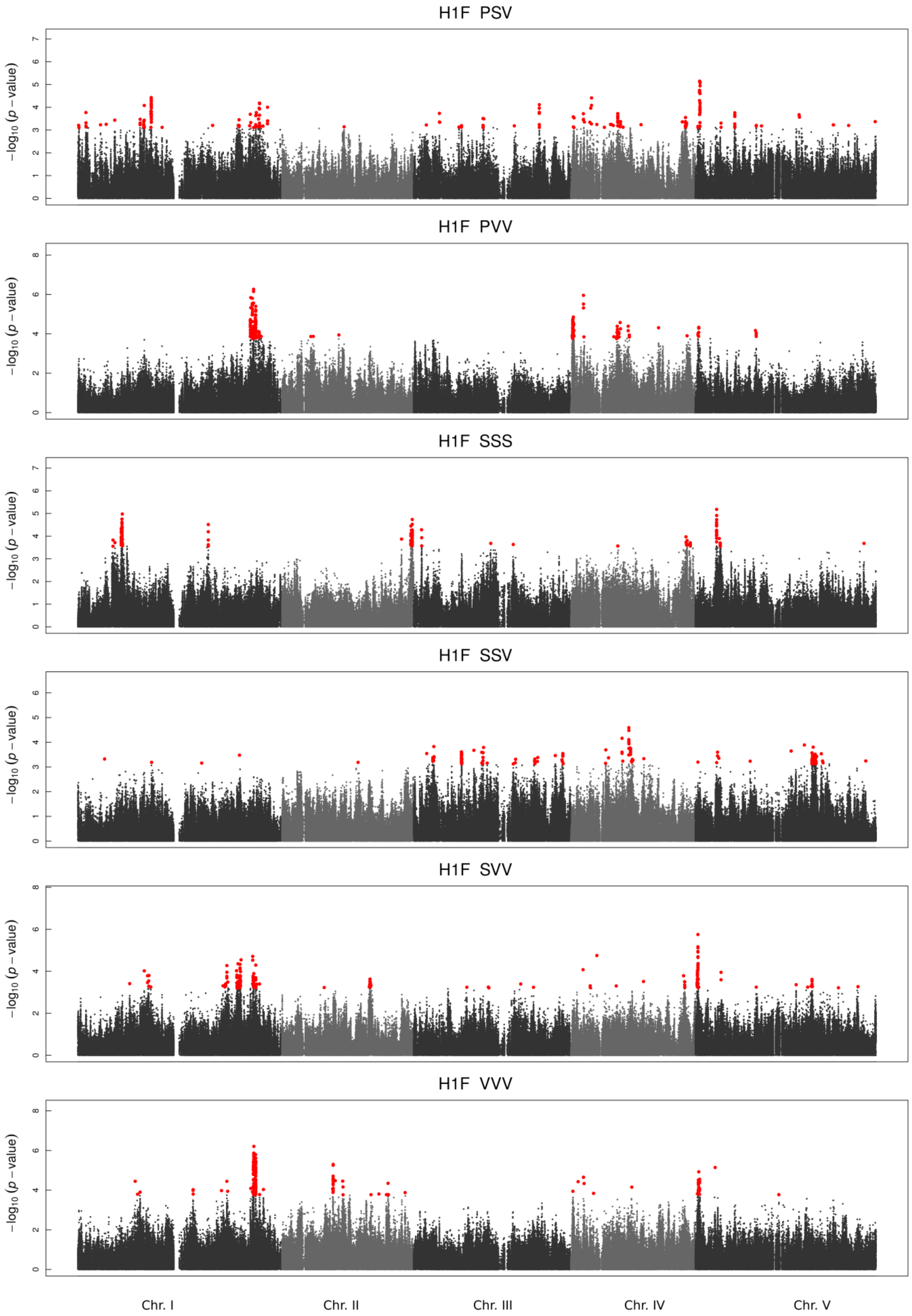

**Figure S2 (continued)**

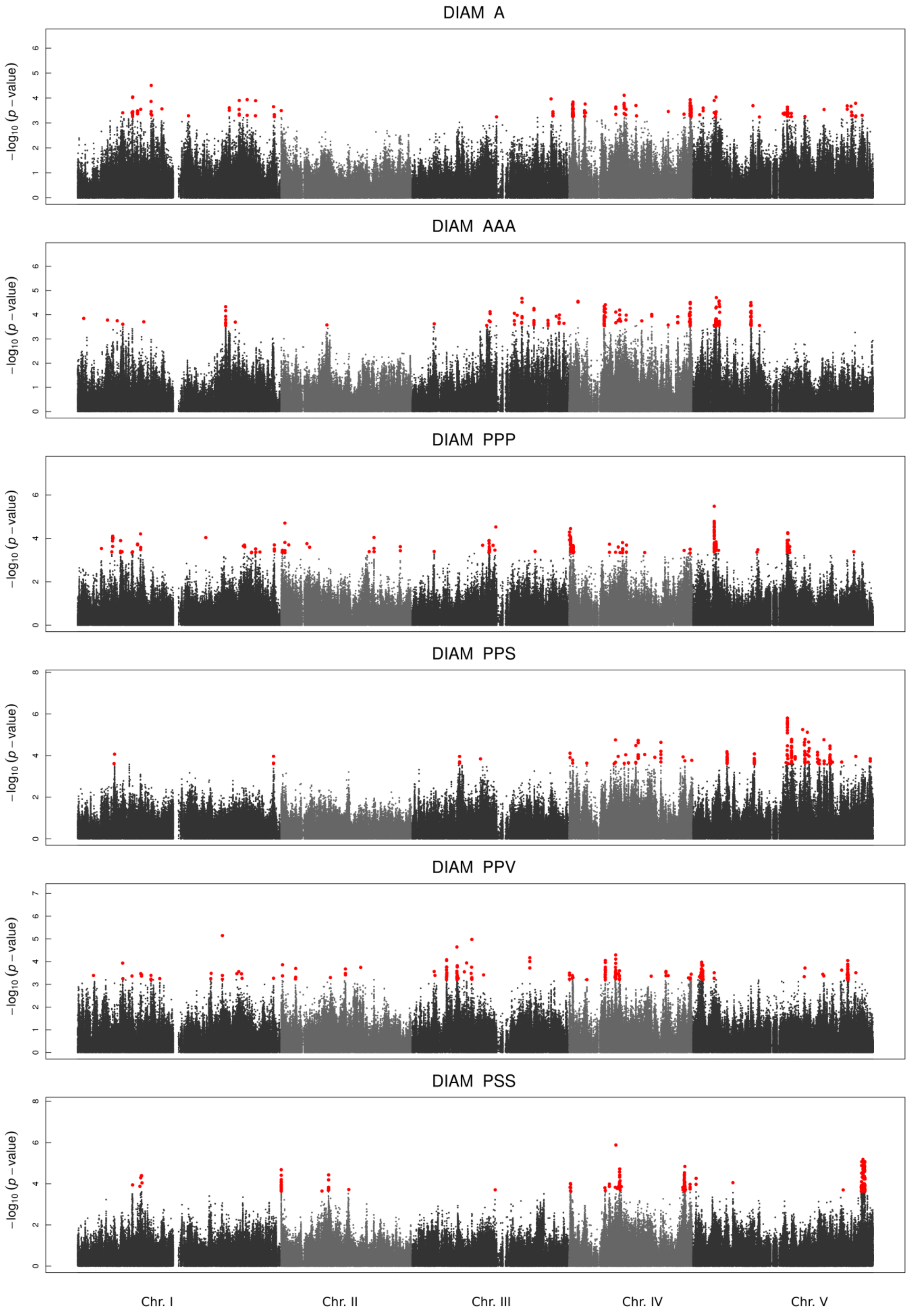

**Figure S2 (continued)**

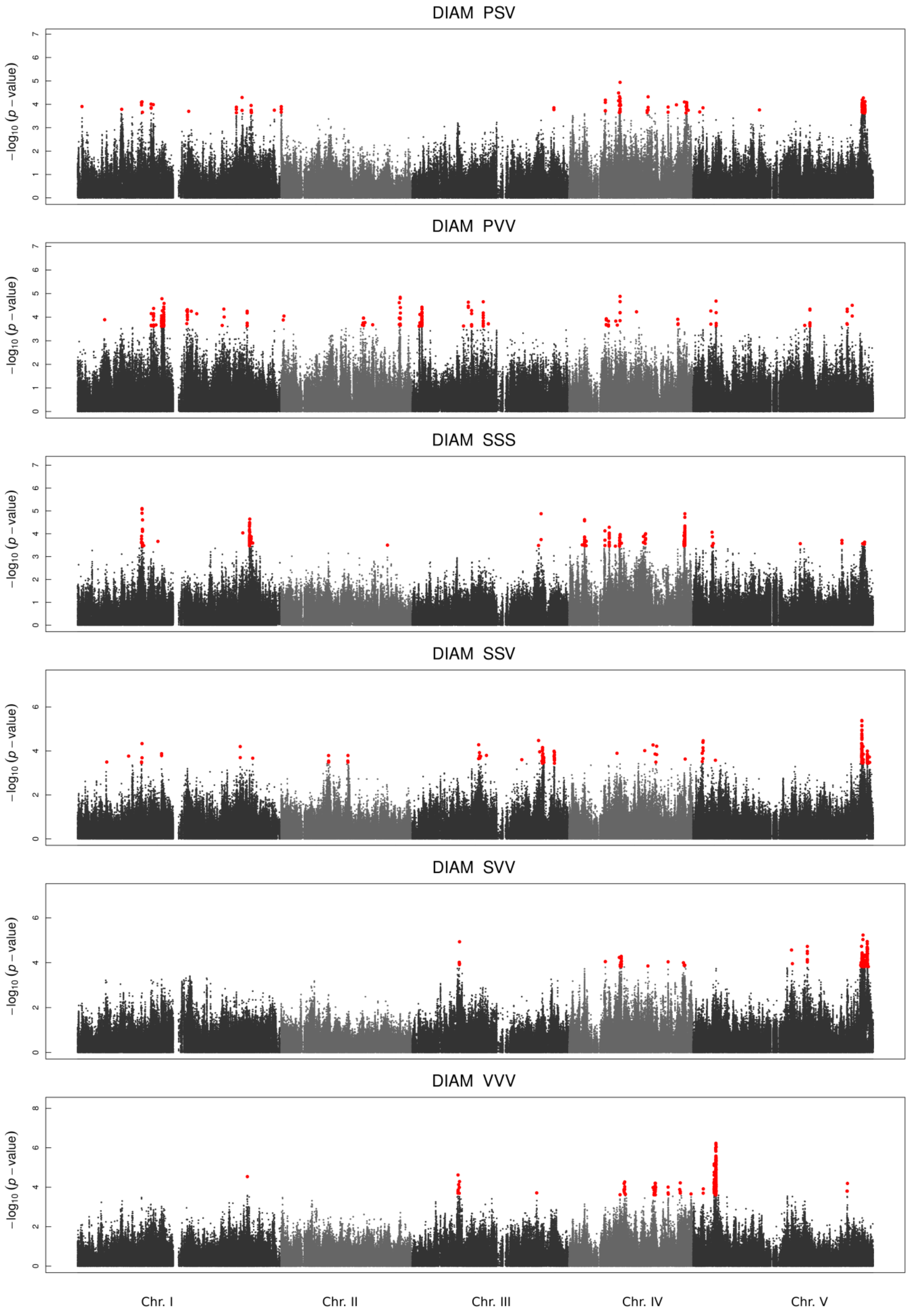

**Figure S2 (continued)**

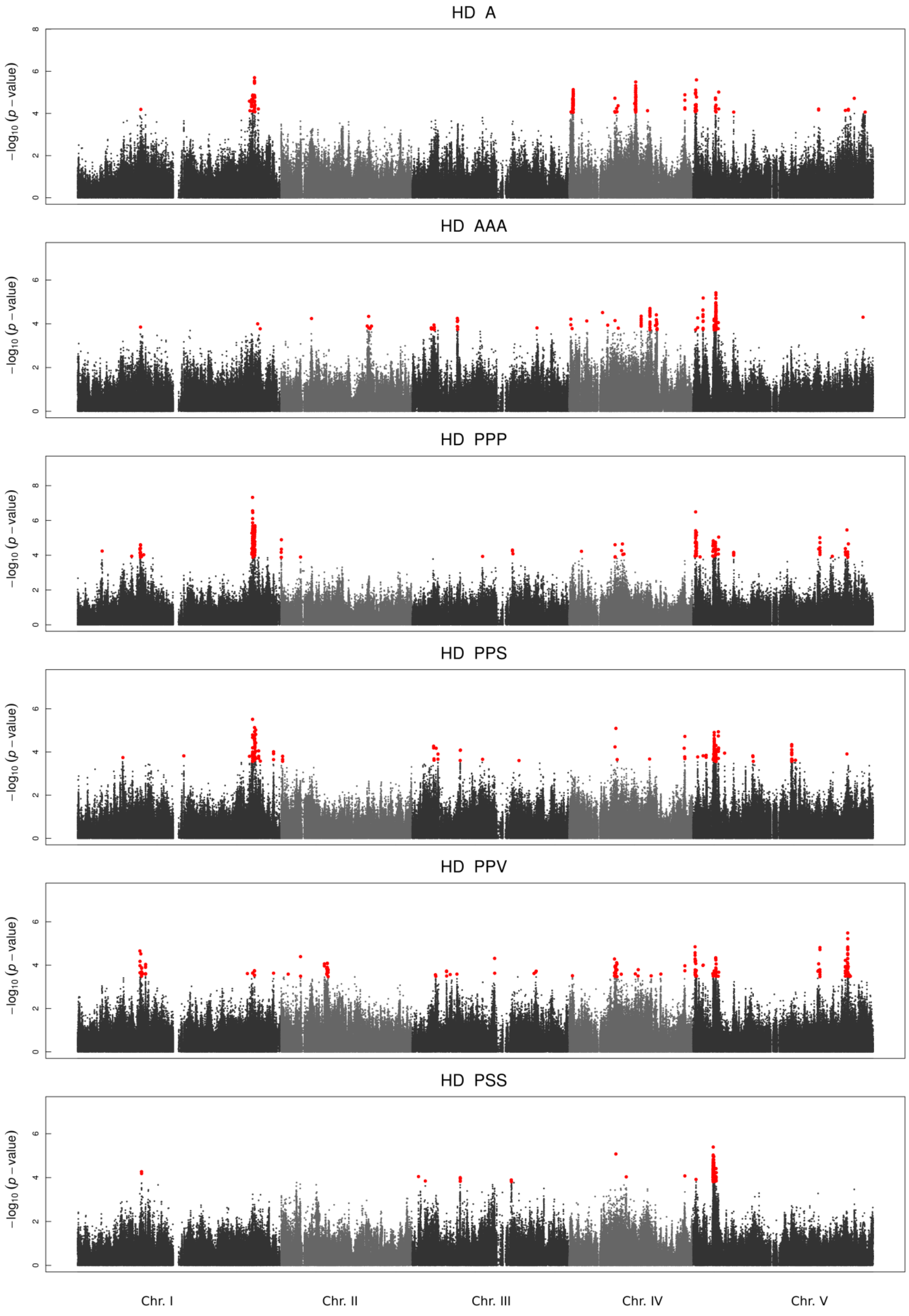

**Figure S2 (continued)**

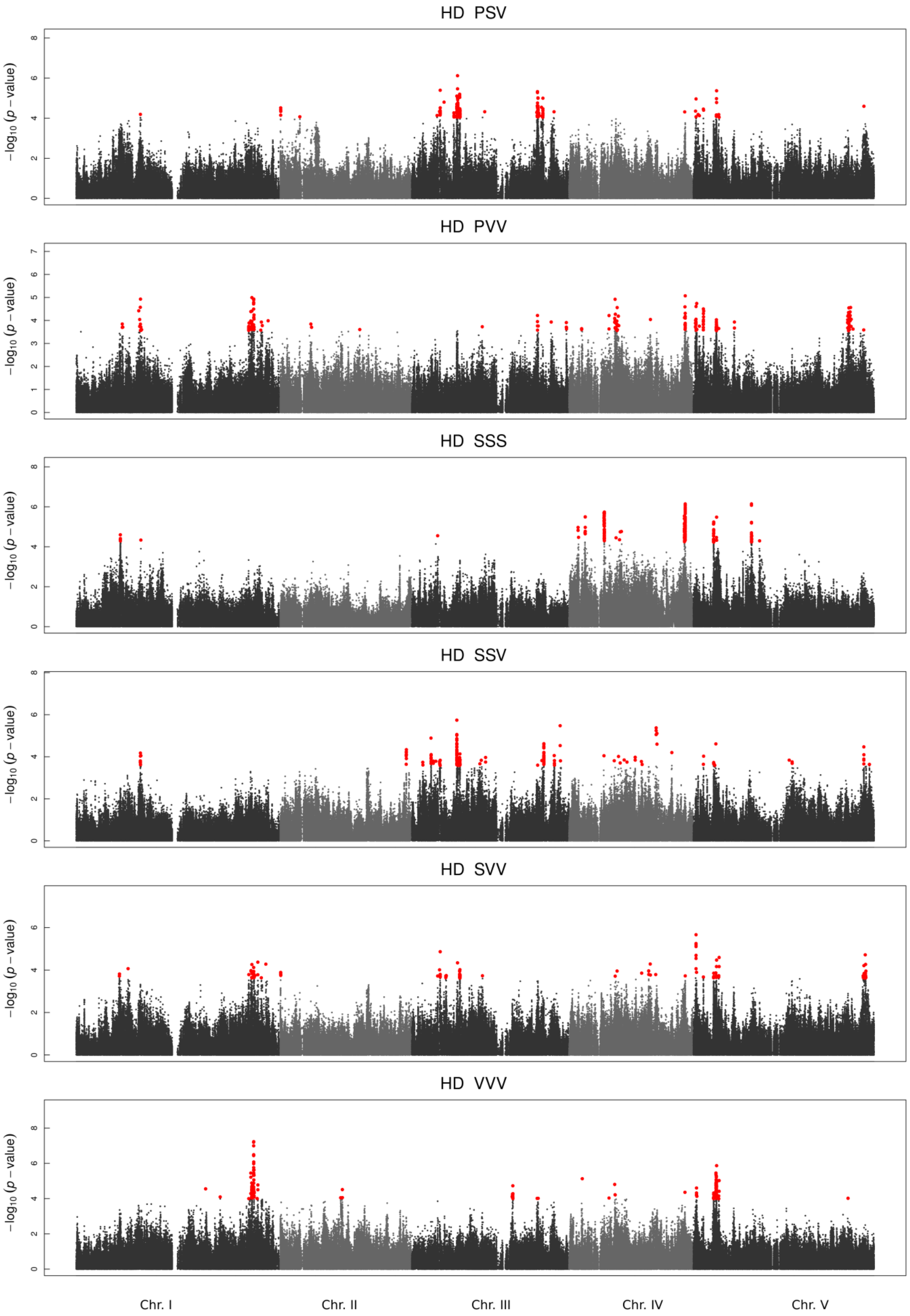

**Figure S2 (continued)**

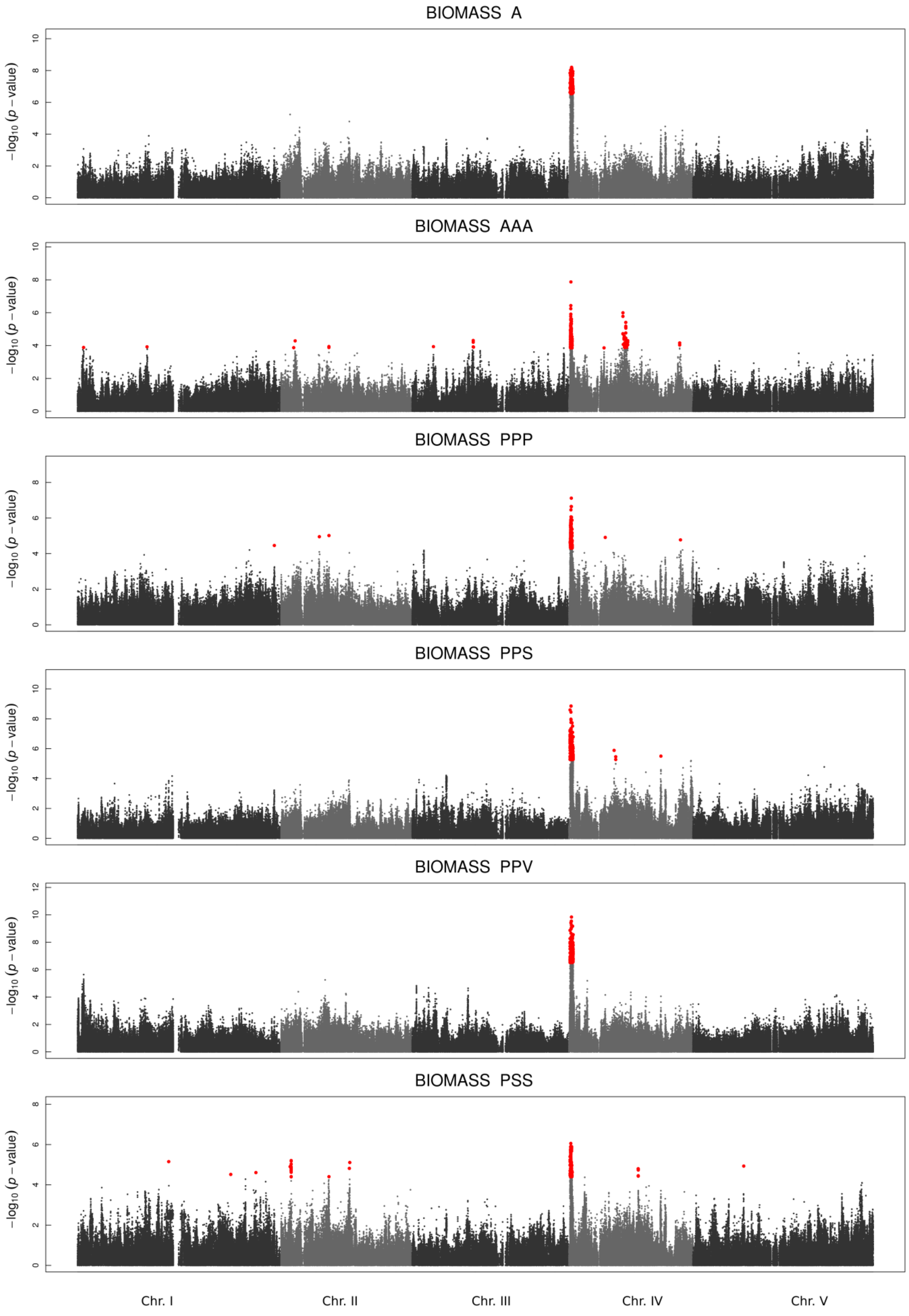

**Figure S2 (continued)**

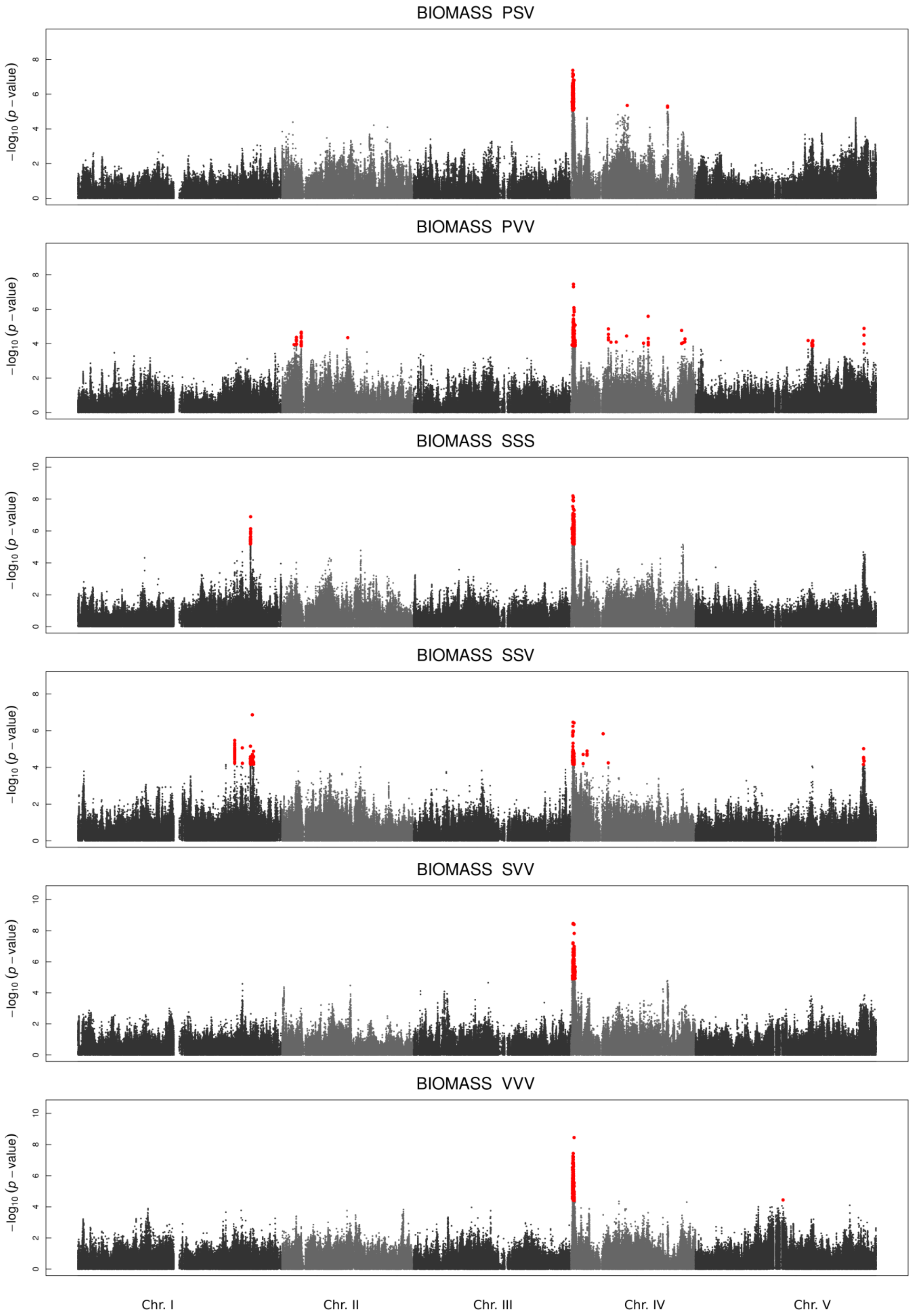

**Figure S3. Non-proportional Venn diagram presenting the partitioning of H1F SNPs detected among the lists of 200 top SNPs for each treatment, according different subsets of treatments.**

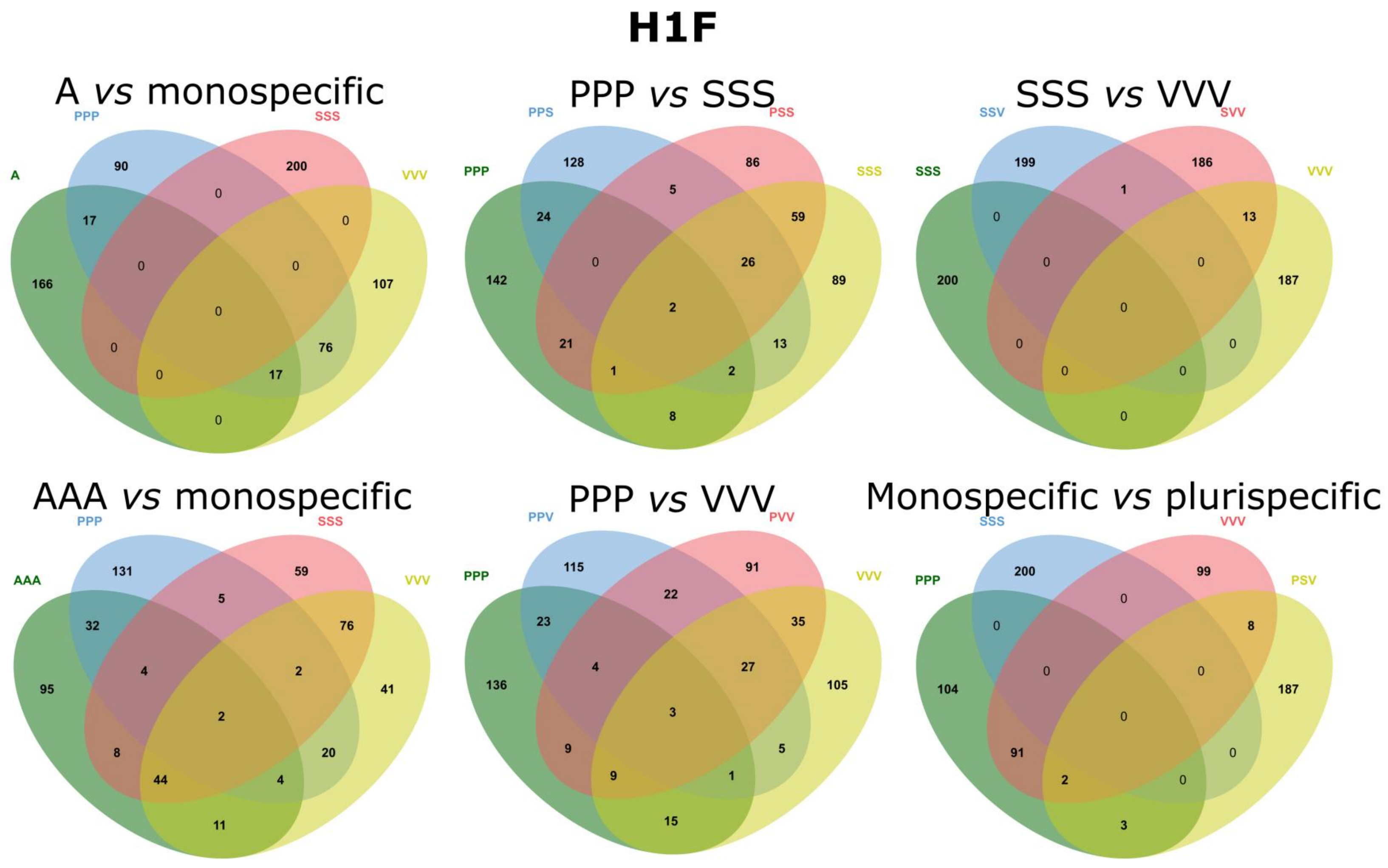

**Figure S4. Non-proportional Venn diagram presenting the partitioning of DIAM SNPs detected among the lists of 200 top SNPs for each treatment, according different subsets of treatments.**

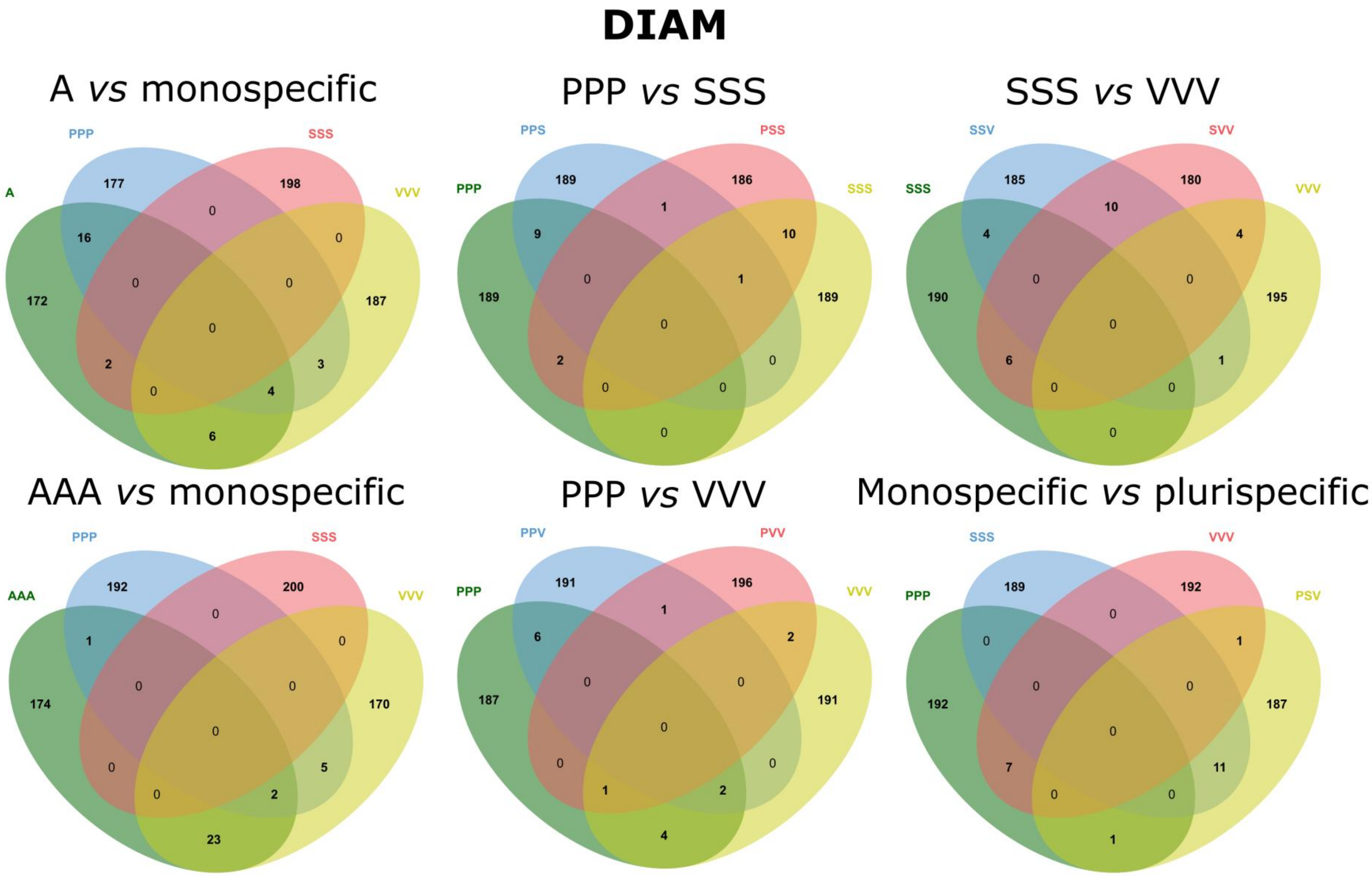

**Figure S5. Non-proportional Venn diagram presenting the partitioning of HD SNPs detected among the lists of 200 top SNPs for each treatment, according different subsets of treatments.**

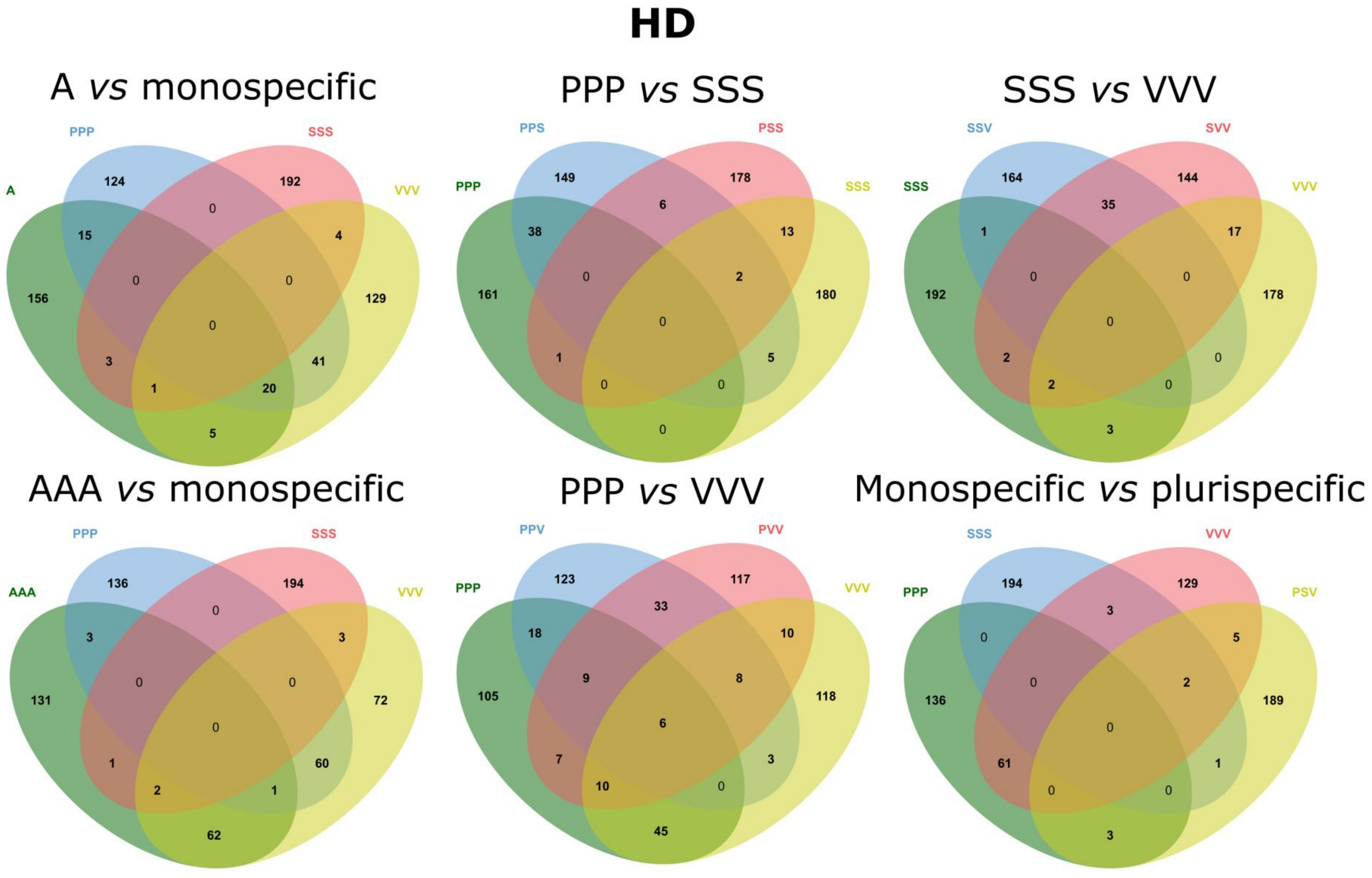

**Figure S6. Non-proportional Venn diagram presenting the partitioning of BIOMASS SNPs detected among the lists of 200 top SNPs for each treatment, according different subsets of treatments.**

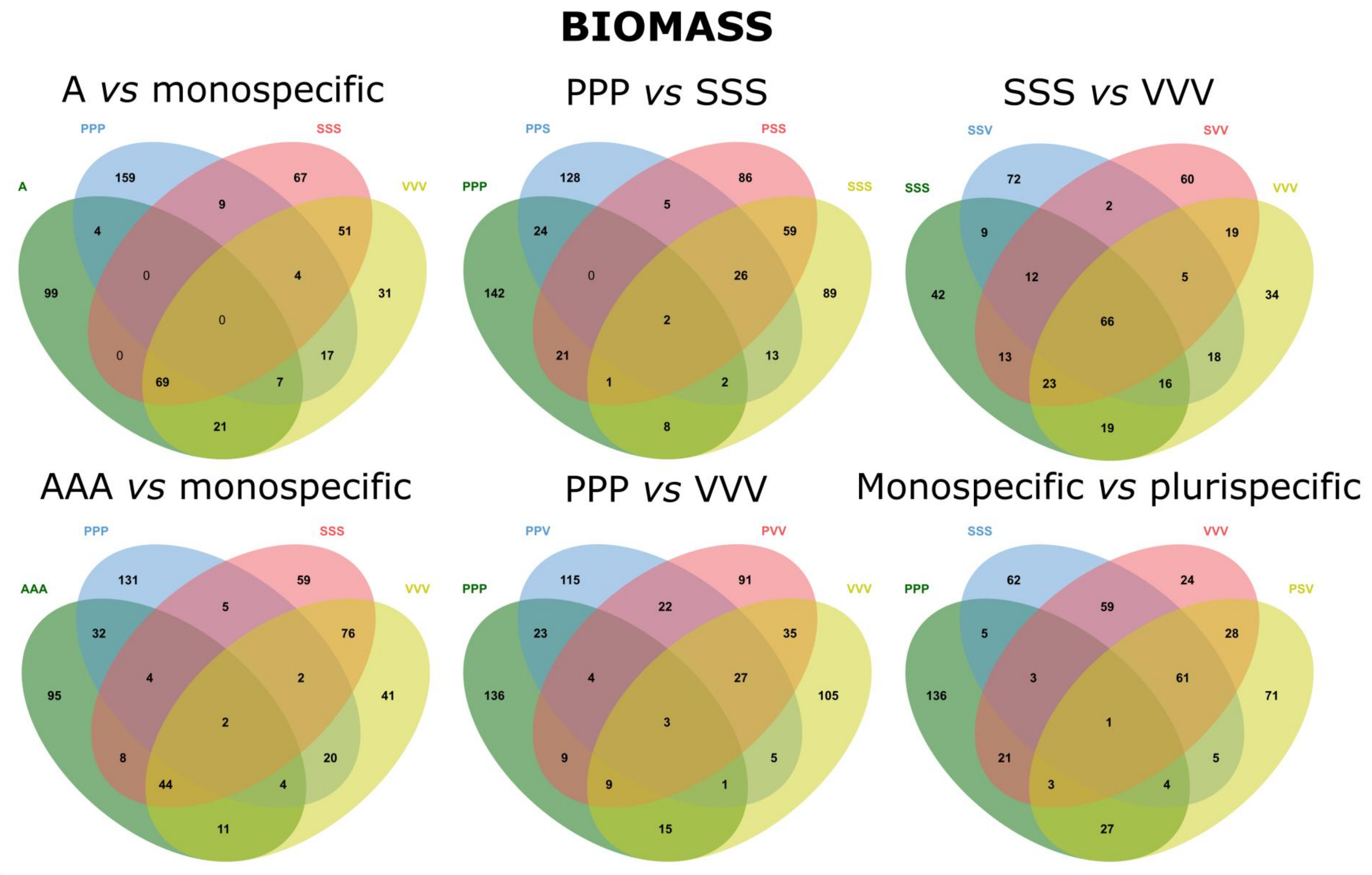

**Table S1 Natural variation of four phenotypic traits scored on *A. thaliana* plants among all treatments, with the exception of the 'Control' treatment.** Bold *P*-values indicate significant effects after FDR correction. Model random terms were tested with likelihood ratio tests (LRT) of models with and without these effects. Random effects are in italics. H1F: height from the soil to the first flower on the main stem, DIAM: maximum diameter of the rosette, BIOMASS: aboveground dry biomass, HD = H1F / DIAM.

| | **Traits** | | | | | | | |
|---|---|---|---|---|---|---|---|---|
| | **H1F** | | **DIAM** | | **BIOMASS** | | **HD** | |
| **Model terms** | *F or LRT* | *P* | *F or LRT* | *P* | *F or LRT* | *P* | *F or LRT* | *P* |
| block | 2.97 | **6.01E-02** | 1.11 | 4.06E-01 | 3.36 | **4.24E-02** | 0.52 | 6.71E-01 |
| treatment | 22.58 | **2.55E-32** | 3.36 | **3.44E-04** | 22.89 | **2.55E-32** | 1.66 | 1.03E-01 |
| *accession* | 822.20 | **1.13E-179** | 718.50 | **2.65E-157** | 450.30 | **4.35E-99** | 984.50 | **1.18E-214** |
| *treatment*accession* | 19.10 | **1.93E-05** | 26.90 | **3.52E-07** | 171.50 | **1.95E-38** | 28.00 | **2.12E-07** |
| germ(treatment) | 1.31 | 2.50E-01 | 0.90 | 5.86E-01 | 2.04 | **3.00E-02** | 0.85 | 6.09E-01 |
| flo(treatment) | 49.74 | **2.55E-32** | 9.30 | **3.42E-16** | 43.33 | **2.55E-32** | 9.65 | **6.94E-17** |
| flo*flo(treatment) | 38.79 | **2.55E-32** | 8.25 | **4.52E-14** | 17.41 | **2.55E-32** | 7.72 | **5.30E-13** |

**Table S2 Natural variation of four phenotypic traits scored on *A. thaliana* plants among all treatments, with the exception of the 'Control' and 'Intraspecific interaction' treatments.** Bold *P*-values indicate significant effects after FDR correction. Model random terms were tested with likelihood ratio tests (LRT) of models with and without these effects. Random effects are in italics. H1F: height from the soil to the first flower on the main stem, DIAM: maximum diameter of the rosette, BIOMASS: aboveground dry biomass, HD = H1F / DIAM.

| | **Traits** | | | | | | | |
|---|---|---|---|---|---|---|---|---|
| | **H1F** | | **DIAM** | | **BIOMASS** | | **HD** | |
| **Model terms** | *F or LRT* | *P* | *F or LRT* | *P* | *F or LRT* | *P* | *F or LRT* | *P* |
| block | 2.95 | **6.39E-02** | 1.32 | 3.23E-01 | 3.80 | **2.89E-02** | 0.44 | 7.28E-01 |
| treatment | 25.42 | **2.80E-32** | 3.38 | **6.14E-04** | 23.94 | **2.80E-32** | 1.84 | 6.87E-02 |
| *accession* | 762.00 | **1.38E-166** | 615.00 | **8.54E-135** | 390.00 | **5.80E-86** | 879.50 | **7.87E-192** |
| *treatment*accession* | 8.70 | **4.69E-03** | 25.60 | **7.35E-07** | 155.60 | **5.82E-35** | 24.70 | **1.10E-06** |
| germ(treatment) | 1.35 | 2.29E-01 | 0.89 | 5.62E-01 | 2.10 | **2.89E-02** | 0.92 | 5.51E-01 |
| flo(treatment) | 53.64 | **2.80E-32** | 9.56 | **2.16E-15** | 44.56 | **2.80E-32** | 10.73 | **1.42E-17** |
| flo*flo(treatment) | 40.04 | **2.80E-32** | 8.88 | **3.66E-14** | 17.48 | **1.02E-30** | 8.54 | **1.55E-13** |

**Table S3 Natural variation of four phenotypic traits scored on *A. thaliana* plants within each treatment.** Bold *P*-values indicate significant effects after FDR correction. Model random terms were tested with likelihood ratio tests (LRT) of models with and without these effects. Random effects are in italics. H1F: height from the soil to the first flower on the main stem, DIAM: maximum diameter of the rosette, BIOMASS: aboveground dry biomass, HD = H1F / DIAM.

| | | H1F | | DIAM | | BIOMASS | | HD | |
|---|---|---|---|---|---|---|---|---|---|
| **Treatment** | **Model Term** | ***F or LRT*** | ***P*** | ***F or LRT*** | ***P*** | ***F or LRT*** | ***P*** | ***F or LRT*** | ***P*** |
| A | block | 2.98 | 5.32E-02 | 0.66 | 6.49E-01 | 19.58 | **9.02E-11** | 4.17 | **1.22E-02** |
| | *accession* | 98.80 | **9.57E-22** | 50.80 | **5.44E-12** | 63.90 | **1.21E-14** | 78.00 | **1.44E-17** |
| | germ | 0.00 | 9.89E-01 | 1.51 | 2.93E-01 | 3.26 | 1.10E-01 | 0.04 | 8.46E-01 |
| | flo | 1.86 | 2.39E-01 | 179.53 | **1.60E-29** | 127.02 | **1.42E-22** | 13.71 | **5.59E-04** |
| | flo*flo | 0.72 | 4.78E-01 | 93.91 | **6.62E-18** | 32.65 | **9.79E-08** | 8.39 | **7.98E-03** |
| AAA | block | 0.45 | 7.63E-01 | 1.59 | 2.62E-01 | 1.33 | 3.49E-01 | 2.17 | 1.39E-01 |
| | *accession* | 70.70 | **4.75E-16** | 41.20 | **5.30E-10** | 14.80 | **2.74E-04** | 42.20 | **3.47E-10** |
| | germ | 5.04 | **4.37E-02** | 0.55 | 5.41E-01 | 0.17 | 7.31E-01 | 3.51 | 9.81E-02 |
| | flo | 0.85 | 4.45E-01 | 109.99 | **2.39E-20** | 100.12 | **7.66E-19** | 22.20 | **1.09E-05** |
| | flo*flo | 0.82 | 4.52E-01 | 87.52 | **4.84E-17** | 48.09 | **1.45E-10** | 16.42 | **1.59E-04** |
| PPP | block | 0.39 | 7.97E-01 | 2.32 | 1.15E-01 | 4.31 | **1.04E-02** | 1.15 | 4.21E-01 |
| | *accession* | 72.50 | **2.00E-16** | 56.80 | **3.62E-13** | 36.10 | **6.70E-09** | 64.50 | **9.26E-15** |
| | germ | 0.95 | 4.22E-01 | 1.64 | 2.72E-01 | 1.95 | 2.28E-01 | 2.61 | 1.59E-01 |
| | flo | 0.23 | 6.89E-01 | 115.96 | **4.72E-21** | 25.93 | **1.93E-06** | 14.27 | **4.38E-04** |
| | flo*flo | 0.05 | 8.46E-01 | 60.27 | **1.36E-12** | 0.26 | 6.81E-01 | 8.33 | **8.23E-03** |
| PPS | block | 3.35 | **3.45E-02** | 1.20 | 3.99E-01 | 7.26 | **2.52E-04** | 3.00 | 5.22E-02 |
| | *accession* | 41.40 | **4.92E-10** | 37.70 | **3.09E-09** | 16.40 | **1.26E-04** | 43.90 | **1.57E-10** |
| | germ | 0.63 | 5.10E-01 | 0.24 | 6.89E-01 | 1.17 | 3.67E-01 | 0.33 | 6.37E-01 |
| | flo | 2.12 | 2.08E-01 | 72.75 | **1.25E-14** | 68.30 | **6.54E-14** | 0.72 | 4.78E-01 |
| | flo*flo | 10.06 | **3.51E-03** | 91.57 | **1.44E-17** | 57.89 | **3.18E-12** | 0.19 | 7.25E-01 |

**Table S3 (continued)**

| | | H1F | | DIAM | | BIOMASS | | HD | |
|---|---|---|---|---|---|---|---|---|---|
| **Treatment** | **Model Term** | ***F or LRT*** | ***P*** | ***F or LRT*** | ***P*** | ***F or LRT*** | ***P*** | ***F or LRT*** | ***P*** |
| PPV | block | 1.08 | 4.45E-01 | 2.44 | 1.01E-01 | 7.65 | **1.59E-04** | 0.82 | 5.62E-01 |
| | *accession* | 66.30 | **3.87E-15** | 21.10 | **1.18E-05** | 61.20 | **4.27E-14** | 67.90 | **1.88E-15** |
| | germ | 0.75 | 4.75E-01 | 0.45 | 5.83E-01 | 4.30 | 6.30E-02 | 0.16 | 7.40E-01 |
| | flo | 5.01 | **4.39E-02** | 229.16 | **1.20E-30** | 96.16 | **4.49E-18** | 10.83 | **2.40E-03** |
| | flo*flo | 4.34 | 6.21E-02 | 147.03 | **5.91E-25** | 31.82 | **1.48E-07** | 6.13 | **2.49E-02** |
| PSS | block | 4.96 | **4.76E-03** | 1.06 | 4.53E-01 | 2.64 | 7.98E-02 | 6.80 | **4.47E-04** |
| | *accession* | 31.20 | **7.66E-08** | 52.70 | **2.33E-12** | 12.90 | **6.99E-04** | 44.10 | **1.45E-10** |
| | germ | 2.29 | 1.89E-01 | 14.94 | **3.22E-04** | 7.09 | **1.49E-02** | 0.29 | 6.61E-01 |
| | flo | 2.02 | 2.18E-01 | 47.03 | **2.50E-10** | 38.47 | **8.07E-09** | 0.02 | 8.94E-01 |
| | flo*flo | 8.98 | **6.01E-03** | 68.23 | **6.99E-14** | 27.81 | **8.37E-07** | 0.60 | 5.20E-01 |
| PSV | block | 10.25 | **6.10E-06** | 2.88 | 6.03E-02 | 13.62 | **9.57E-08** | 11.30 | **1.62E-06** |
| | *accession* | 31.30 | **7.37E-08** | 46.70 | **4.05E-11** | 9.00 | **5.40E-03** | 52.60 | **2.39E-12** |
| | germ | 0.07 | 8.22E-01 | 1.15 | 3.70E-01 | 2.16 | 2.04E-01 | 0.10 | 7.90E-01 |
| | flo | 2.71 | 1.51E-01 | 55.76 | **7.20E-12** | 61.28 | **9.33E-13** | 0.17 | 7.31E-01 |
| | flo*flo | 10.57 | **2.71E-03** | 63.54 | **4.12E-13** | 45.22 | **4.92E-10** | 2.37 | 1.82E-01 |
| PVV | block | 0.30 | 8.46E-01 | 8.30 | **6.93E-05** | 6.65 | **5.32E-04** | 1.81 | 2.06E-01 |
| | *accession* | 38.40 | **2.19E-09** | 48.90 | **1.35E-11** | 53.10 | **2.00E-12** | 27.00 | **5.87E-07** |
| | germ | 1.10 | 3.82E-01 | 0.06 | 8.30E-01 | 1.86 | 2.39E-01 | 0.73 | 4.78E-01 |
| | flo | 0.73 | 4.78E-01 | 117.67 | **3.36E-21** | 38.65 | **7.55E-09** | 19.31 | **4.21E-05** |
| | flo*flo | 0.43 | 5.91E-01 | 63.36 | **4.42E-13** | 6.77 | **1.76E-02** | 11.14 | **2.05E-03** |

**Table S3 (continued)**

| Treatment | Model Terms | H1F | | DIAM | | BIOMASS | | HD | |
|---|---|---|---|---|---|---|---|---|---|
| | | ***F or LRT*** | ***P*** | ***F or LRT*** | ***P*** | ***F or LRT*** | ***P*** | ***F or LRT*** | ***P*** |
| SSS | block | 2.38 | 1.08E-01 | 7.03 | **3.41E-04** | 9.03 | **2.87E-05** | 5.08 | **4.13E-03** |
| | *accession* | 16.10 | **1.46E-04** | 41.40 | **4.92E-10** | 18.70 | **3.95E-05** | 42.90 | **2.51E-10** |
| | germ | 0.11 | 7.84E-01 | 0.52 | 5.50E-01 | 2.20 | 2.00E-01 | 2.06 | 2.15E-01 |
| | flo | 5.40 | **3.65E-02** | 29.87 | **3.42E-07** | 8.80 | **6.53E-03** | 2.62 | 1.59E-01 |
| | flo*flo | 18.71 | **5.78E-05** | 55.21 | **9.96E-12** | 7.17 | **1.45E-02** | 5.07 | **4.35E-02** |
| SSV | block | 3.51 | **2.82E-02** | 0.70 | 6.28E-01 | 3.26 | **3.83E-02** | 4.19 | **1.20E-02** |
| | *accession* | 7.70 | **1.04E-02** | 28.20 | **3.23E-07** | 37.60 | **3.20E-09** | 15.30 | **2.14E-04** |
| | germ | 1.26 | 3.47E-01 | 0.11 | 7.84E-01 | 0.33 | 6.37E-01 | 2.48 | 1.71E-01 |
| | flo | 0.85 | 4.45E-01 | 45.10 | **4.92E-10** | 21.37 | **1.62E-05** | 4.40 | 6.04E-02 |
| | flo*flo | 7.71 | **1.11E-02** | 61.94 | **7.34E-13** | 16.22 | **1.76E-04** | 0.60 | 5.20E-01 |
| SVV | block | 6.81 | **4.38E-04** | 2.88 | 6.03E-02 | 13.37 | **1.26E-07** | 9.29 | **2.02E-05** |
| | *accession* | 33.40 | **2.54E-08** | 29.60 | **1.63E-07** | 18.30 | **4.77E-05** | 28.60 | **2.67E-07** |
| | germ | 0.00 | 9.89E-01 | 0.88 | 4.40E-01 | 3.40 | 1.03E-01 | 0.10 | 7.90E-01 |
| | flo | 1.48 | 2.98E-01 | 91.53 | **1.44E-17** | 77.54 | **2.23E-15** | 3.45 | 1.01E-01 |
| | flo*flo | 7.59 | **1.18E-02** | 96.48 | **3.10E-18** | 56.58 | **5.18E-12** | 0.36 | 6.26E-01 |
| VVV | block | 0.75 | 5.98E-01 | 4.32 | **1.04E-02** | 2.56 | 8.79E-02 | 0.51 | 7.31E-01 |
| | *accession* | 51.60 | **3.80E-12** | 36.30 | **6.15E-09** | 61.70 | **3.43E-14** | 53.00 | **2.06E-12** |
| | germ | 1.70 | 2.62E-01 | 0.05 | 8.46E-01 | 1.53 | 2.91E-01 | 2.75 | 1.48E-01 |
| | flo | 0.00 | 9.76E-01 | 195.71 | **1.20E-30** | 100.86 | **7.82E-19** | 36.83 | **1.63E-08** |
| | flo*flo | 0.25 | 6.86E-01 | 129.57 | **8.50E-23** | 46.31 | **3.23E-10** | 30.89 | **2.12E-07** |

**Table S4 Broad-sense heritability values for the four phenotypic traits scored on *A. thaliana* plants within each treatment.** Bold *P*-values indicate significant effects after FDR correction.

| | Traits | | | | | | | |
|---|---|---|---|---|---|---|---|---|
| | H1F | | DIAM | | BIOMASS | | HD | |
| Treatment | $H^2$ | *P* | $H^2$ | *P* | $H^2$ | *P* | $H^2$ | *P* |
| A | 0.8 | **9.57E-22** | 0.94 | **5.44E-12** | 0.96 | **1.21E-14** | 0.86 | **1.44E-17** |
| AAA | 0.74 | **4.75E-16** | 0.89 | **5.30E-10** | 0.91 | **2.74E-04** | 0.82 | **3.47E-10** |
| PPP | 0.77 | **2.00E-16** | 0.95 | **3.62E-13** | 0.96 | **6.70E-09** | 0.89 | **9.26E-15** |
| SSS | 0.8 | **1.46E-04** | 0.67 | **4.92E-10** | 0.65 | **3.95E-05** | 0.8 | **2.51E-10** |
| VVV | 0.68 | **3.80E-12** | 0.94 | **6.15E-09** | 0.94 | **3.43E-14** | 0.85 | **2.06E-12** |
| PPS | 0.82 | **4.92E-10** | 0.71 | **3.09E-09** | 0.69 | **1.26E-04** | 0.84 | **1.57E-10** |
| PPV | 0.75 | **3.87E-15** | 0.93 | **1.18E-05** | 0.96 | **4.27E-14** | 0.89 | **1.88E-15** |
| PSS | 0.81 | **7.66E-08** | 0.75 | **2.33E-12** | 0.74 | **6.99E-04** | 0.82 | **1.45E-10** |
| PVV | 0.65 | **2.19E-09** | 0.94 | **1.35E-11** | 0.94 | **2.00E-12** | 0.83 | **5.87E-07** |
| SSV | 0.77 | **1.04E-02** | 0.73 | **3.23E-07** | 0.78 | **3.20E-09** | 0.78 | **2.14E-04** |
| SVV | 0.81 | **2.54E-08** | 0.76 | **1.63E-07** | 0.79 | **4.77E-05** | 0.83 | **2.67E-07** |
| PSV | 0.75 | **7.37E-08** | 0.77 | **4.05E-11** | 0.69 | **5.40E-03** | 0.81 | **2.39E-12** |

**Table S5 Biological pathways (MapMan classification) represented for all unique genes identified.** Bold lines indicate significant over-represented biological pathways ($P < 0.01$).

| Normed Freq. | ± bootstrap StdDev | p-value | Class |
|---|---|---|---|
| **0.17** | **0.078** | **7.96E-10** | **DNA** |
| **1.87** | **0.425** | **2.53E-03** | **transport** |
| **0.86** | **0.061** | **6.75E-03** | **not assigned** |
| 1.25 | 0.166 | 0.021 | RNA |
| 1.4 | 0.313 | 0.029 | signalling |
| 1.14 | 0.142 | 0.029 | protein |
| 3.45 | 1.741 | 0.045 | TCA / org transformation |
| 1.69 | 0.665 | 0.054 | lipid metabolism |
| 4.55 | 3.366 | 0.062 | C1-metabolism |
| 1.2 | 0.216 | 0.063 | misc |
| 3.79 | 2.729 | 0.082 | tetrapyrrole synthesis |
| 0.81 | 0.237 | 0.094 | stress |
| 2.2 | 1.051 | 0.108 | minor CHO metabolism |
| 1.76 | 0.859 | 0.114 | PS |
| 0.87 | 0.31 | 0.132 | cell |
| 1.04 | 0.352 | 0.133 | development |
| 1.16 | 0.418 | 0.14 | cell wall |
| 1 | 0.415 | 0.163 | hormone metabolism |
| 1.02 | 0.435 | 0.177 | secondary metabolism |
| 0.98 | 0.363 | 0.178 | micro RNA, natural antisense etc |
| 1.8 | 1.331 | 0.204 | major CHO metabolism |
| 1.3 | 0.676 | 0.206 | redox |
| 3.5 | 2.423 | 0.217 | N-metabolism |
| 3.25 | 2.753 | 0.228 | Biodegradation of Xenobiotics |
| 0.7 | 0.458 | 0.236 | amino acid metabolism |
| 0.5 | 0.4 | 0.274 | nucleotide metabolism |
| 0.6 | 0.44 | 0.317 | mitochondrial electron transport / ATP synthesis |
| 1.15 | 0.907 | 0.367 | glycolysis |
| 1.12 | 0.622 | 0.368 | Co-factor and vitamine metabolism |
| 1.09 | 0.846 | 0.369 | metal handling |

**Table S6 Biological pathways (MapMan classification) represented in the 'Control', 'Intraspecific', 'Monospecific' and 'Plurispecific' treatments.** Bold lines indicate significant over-represented biological pathways ($P < 0.01$).

| Type | Absolute values | ± bootstrap | p-value | Class |
|---|---|---|---|---|
| Control | 10 | 2.8 | 0.02 | not assigned |
| Control | 2 | 1.2 | 0.03 | PS |
| Control | 1 | 0.8 | 0.052 | C1-metabolism |
| Control | 1 | 0.8 | 0.062 | tetrapyrrole synthesis |
| Control | 3 | 1.5 | 0.071 | development |
| Control | 1 | 0.7 | 0.097 | TCA / org transformation |
| Control | 4 | 1.9 | 0.107 | misc |
| Control | 6 | 2.4 | 0.116 | RNA |
| Control | 8 | 2.5 | 0.131 | protein |
| Control | 1 | 0.6 | 0.144 | minor CHO metabolism |
| Control | 3 | 1.4 | 0.15 | stress |
| Control | 2 | 1.1 | 0.246 | transport |
| Control | 2 | 1.2 | 0.276 | signalling |
| Control | 1 | 0.9 | 0.33 | lipid metabolism |
| Control | 1 | 0.7 | 0.339 | micro RNA, natural antisense etc |
| Control | 1 | 0.8 | 0.358 | cell wall |
| Intraspecific | 2 | 1.2 | 0.092 | DNA |
| Intraspecific | 1 | 0.7 | 0.107 | TCA / org transformation |
| Intraspecific | 18 | 2.8 | 0.116 | not assigned |
| Intraspecific | 2 | 1.3 | 0.116 | secondary metabolism |
| Intraspecific | 1 | 1 | 0.132 | major CHO metabolism |
| Intraspecific | 6 | 2.1 | 0.139 | RNA |
| Intraspecific | 8 | 2.6 | 0.147 | protein |
| Intraspecific | 3 | 1.4 | 0.191 | signalling |
| Intraspecific | 1 | 0.7 | 0.214 | misc |
| Intraspecific | 2 | 1.1 | 0.219 | development |

**Table S6 (continued)**

| Type | Absolute values | ± bootstrap | p-value | Class |
|---|---|---|---|---|
| Intraspecific | 2 | 1.1 | 0.26 | transport |
| Intraspecific | 1 | 0.8 | 0.268 | amino acid metabolism |
| Intraspecific | 1 | 0.7 | 0.343 | lipid metabolism |
| Intraspecific | 1 | 0.7 | 0.361 | cell |
| Intraspecific | 1 | NaN | 0.365 | hormone metabolism |
| Intraspecific | 1 | 0.7 | 0.366 | cell wall |
| **Monospecific** | **2** | **1.2** | **8.33E-03** | **tetrapyrrole synthesis** |
| Monospecific | 25 | 4.3 | 0.011 | not assigned |
| Monospecific | 2 | 1.1 | 0.021 | TCA / org transformation |
| Monospecific | 14 | 3 | 0.028 | RNA |
| Monospecific | 17 | 4.2 | 0.076 | protein |
| Monospecific | 2 | 1.1 | 0.099 | PS |
| Monospecific | 3 | 1.7 | 0.105 | micro RNA, natural antisense etc |
| Monospecific | 3 | 1.4 | 0.136 | hormone metabolism |
| Monospecific | 5 | 2.1 | 0.139 | stress |
| Monospecific | 5 | 2.2 | 0.159 | signalling |
| Monospecific | 4 | 2 | 0.19 | misc |
| Monospecific | 3 | 1.5 | 0.206 | development |
| Monospecific | 3 | 1.4 | 0.213 | cell |
| Monospecific | 2 | 1.1 | 0.226 | lipid metabolism |
| Monospecific | 3 | 1.8 | 0.228 | transport |
| Monospecific | 2 | 1 | 0.232 | secondary metabolism |
| Monospecific | 1 | 0.9 | 0.253 | minor CHO metabolism |
| Monospecific | 1 | 0.6 | 0.324 | cell wall |
| Monospecific | 1 | 0.6 | 0.334 | redox |

**Table S6 (continued)**

| Type | Absolute values | ± bootstrap | p-value | Class |
|---|---|---|---|---|
| **Plurispecific** | **5** | **2** | **1.24E-06** | **DNA** |
| **Plurispecific** | **16** | **3.7** | **3.26E-03** | **transport** |
| **Plurispecific** | **18** | **4** | **8.90E-03** | **signalling** |
| Plurispecific | 78 | 7.5 | 0.014 | not assigned |
| Plurispecific | 31 | 5 | 0.021 | RNA |
| Plurispecific | 2 | 1.3 | 0.034 | C1-metabolism |
| Plurispecific | 40 | 6.2 | 0.06 | protein |
| Plurispecific | 15 | 3.5 | 0.076 | misc |
| Plurispecific | 2 | 1.2 | 0.1 | TCA / org transformation |
| Plurispecific | 1 | 0.7 | 0.103 | micro RNA, natural antisense etc |
| Plurispecific | 8 | 3.1 | 0.127 | stress |
| Plurispecific | 4 | 1.8 | 0.133 | development |
| Plurispecific | 3 | 1.6 | 0.139 | redox |
| Plurispecific | 6 | 2.2 | 0.161 | cell |
| Plurispecific | 1 | 0.9 | 0.164 | N-metabolism |
| Plurispecific | 5 | 2 | 0.165 | cell wall |
| Plurispecific | 1 | 0.9 | 0.174 | Biodegradation of Xenobiotics |
| Plurispecific | 2 | 1.1 | 0.175 | minor CHO metabolism |
| Plurispecific | 4 | 1.8 | 0.183 | lipid metabolism |
| Plurispecific | 3 | 1.5 | 0.189 | hormone metabolism |
| Plurispecific | 2 | 1.1 | 0.193 | secondary metabolism |
| Plurispecific | 1 | 0.7 | 0.256 | tetrapyrrole synthesis |
| Plurispecific | 2 | 1.3 | 0.259 | PS |
| Plurispecific | 2 | 1.1 | 0.273 | amino acid metabolism |
| Plurispecific | 1 | 0.9 | 0.333 | glycolysis |

**Table S6 (continued)**

| Type | Absolute values | ± bootstrap | p-value | Class |
|---|---|---|---|---|
| Plurispecific | 1 | 0.6 | 0.336 | Co-factor and vitamine metabolism |
| Plurispecific | 1 | 0.7 | 0.339 | metal handling |
| Plurispecific | 1 | 0.6 | 0.349 | nucleotide metabolism |
| Plurispecific | 1 | 0.7 | 0.359 | major CHO metabolism |
| Plurispecific | 1 | 0.6 | 0.366 | mitochondrial electron transport / ATP synthesis |

### Table S7 Candidate genes underlying the enriched biological pathways (see Table S6).

| Treatment Type | Class | Trait | Treatment | Trait_Treatment | AGI | Subcategory | Annotation |
|---|---|---|---|---|---|---|---|
| Monospecific | tetrapyrrole synthesis | HD | PPP | HD_PPP | AT1G69720 | heme oxygenase | HO3__heme oxygenase 3 |
| | | HD | PPP | HD_PPP | AT1G69740 | ALA dehydratase | HEMB1__Aldolase superfamily protein |
| Plurispecific | DNA | HD | PSV | HD_PSV | AT3G13170 | synthesis/chromatin structure | ATSPO11-1__Spo11/DNA topoisomerase VI, subunit A protein |
| | | HD | SVV | HD_SVV | AT3G13170 | synthesis/chromatin structure | ATSPO11-1__Spo11/DNA topoisomerase VI, subunit A protein |
| | | DIAM | SSV | DIAM_SSV | AT3G50880 | repair | DNA glycosylase superfamily protein |
| | | BIOMASS | PSS | BIOMASS_PSS | AT4G00660 | synthesis/chromatin structure | ATRH8_RH8__RNAhelicase-like 8 |
| | | HD | PPV | HD_PPV | AT5G57160 | synthesis/chromatin structure | ATLIG4_LIG4__DNA ligase IV |
| | | H1F | PSS | H1F_PSS | AT5G59970 | synthesis/chromatin structure histone core H4 | Histone superfamily protein |
| | signalling | DIAM | PSV | DIAM_PSV | AT1G66150 | receptor kinases leucine rich repeat IX | TMK1__transmembrane kinase 1 |
| | | H1F | PVV | H1F_PVV | AT1G69270 | receptor kinases misc | AtRPK1_RPK1__receptor-like protein kinase 1 |
| | | H1F | PSS | H1F_PSS | AT1G69730 | receptor kinases wall associated kinase | Wall-associated kinase family protein |
| | | H1F | PVV | H1F_PVV | AT1G69730 | receptor kinases wall associated kinase | Wall-associated kinase family protein |
| | | HD | PVV | HD_PVV | AT1G69730 | receptor kinases wall associated kinase | Wall-associated kinase family protein |
| | | HD | SVV | HD_SVV | AT1G69730 | receptor kinases wall associated kinase | Wall-associated kinase family protein |
| | | H1F | PPS | H1F_PPS | AT1G70450 | receptor kinases proline extensin like | Protein kinase superfamily protein |
| | | H1F | PVV | H1F_PVV | AT1G70450 | receptor kinases proline extensin like | Protein kinase superfamily protein |
| | | H1F | PPS | H1F_PPS | AT1G70460 | receptor kinases proline extensin like | AtPERK13_PERK13_RHS10__root hair specific 10 |
| | | H1F | PVV | H1F_PVV | AT1G70460 | receptor kinases proline extensin like | AtPERK13_PERK13_RHS10__root hair specific 10 |
| | | DIAM | PVV | DIAM_PVV | AT2G43040 | calcium | NPG1__tetratricopeptide repeat (TPR)-containing protein |
| | | HD | PSV | HD_PSV | AT3G19700 | receptor kinases leucine rich repeat XI | IKU2__Leucine-rich repeat protein kinase family protein |
| | | DIAM | SVV | DIAM_SVV | AT3G20290 | calcium | ATEHD1_EHD1__EPS15 homology domain 1 |
| | | BIOMASS | PSV | BIOMASS_PSV | AT4G00820 | calcium | iqd17__IQ-domain 17 |
| | | BIOMASS | SVV | BIOMASS_SVV | AT4G00820 | calcium | iqd17__IQ-domain 17 |
| | | HD | SVV | HD_SVV | AT4G23160 | receptor kinases DUF 26 | CRK8__cysteine-rich RLK (RECEPTOR-like protein kinase) 8 |
| | | HD | SVV | HD_SVV | AT4G23170 | receptor kinases misc | CRK9_EP1__receptor-like protein kinase-related family protein |
| | | HD | PVV | HD_PVV | AT5G05160 | receptor kinases leucine rich repeat III | RUL1__Leucine-rich repeat protein kinase family protein |
| | | H1F | PSS | H1F_PSS | AT5G10290 | receptor kinases leucine rich repeat II | leucine-rich repeat transmembrane protein kinase family protein |
| | | H1F | PSS | H1F_PSS | AT5G10450 | 14-3-3 proteins | 14-3-3lambda_AFT1_GRF6__G-box regulating factor 6 |
| | | H1F | SSV | H1F_SSV | AT5G40645 | misc | RPM1-interacting protein 4 (RIN4) family protein |
| | | DIAM | PSS | DIAM_PSS | AT5G63710 | receptor kinases leucine rich repeat II | Leucine-rich repeat protein kinase family protein |
| | | HD | SVV | HD_SVV | AT5G64260 | in sugar and nutrient physiology | EXL2__EXORDIUM like 2 |
| | | DIAM | SVV | DIAM_SVV | AT5G65240 | receptor kinases leucine rich repeat II | Leucine-rich repeat protein kinase family protein |
| | transport | H1F | PSV | H1F_PSV | AT1G30840 | nucleotides | ATPUP4_PUP4__purine permease 4 |
| | | H1F | PVV | H1F_PVV | AT1G68740 | phosphate | PHO1;H1__EXS (ERD1/XPR1/SYG1) family protein |
| | | H1F | PSV | H1F_PSV | AT1G72125 | peptides and oligopeptides | AtNPF5.13_NPF5.13__Major facilitator superfamily protein |
| | | DIAM | PVV | DIAM_PVV | AT3G05160 | sugars | Major facilitator superfamily protein |
| | | HD | PSV | HD_PSV | AT3G19640 | unspecified cations | MGT4_MRS2-3__magnesium transporter 4 |
| | | HD | PSV | HD_PSV | AT3G20460 | sugars | Major facilitator superfamily protein |
| | | DIAM | PVV | DIAM_PVV | AT3G23430 | phosphate | ATPHO1_PHO1__phosphate 1 |
| | | DIAM | PVV | DIAM_PVV | AT3G28390 | ABC transporters and multidrug resistance systems | ABCB18_PGP18__P-glycoprotein 18 |
| | | DIAM | PPV | DIAM_PPV | AT3G47740 | ABC transporters and multidrug resistance systems | ABCA3_ATATH2_ATH2__ABC2 homolog 2 |
| | | BIOMASS | PPV | BIOMASS_PPV | AT4G00900 | calcium | ATECA2_ECA2__ER-type Ca2+-ATPase 2 |
| | | BIOMASS | PSS | BIOMASS_PSS | AT4G00900 | calcium | ATECA2_ECA2__ER-type Ca2+-ATPase 2 |
| | | BIOMASS | PVV | BIOMASS_PVV | AT4G00900 | calcium | ATECA2_ECA2__ER-type Ca2+-ATPase 2 |
| | | H1F | SSV | H1F_SSV | AT4G15215 | ABC transporters and multidrug resistance systems | ABCG41_ATPDR13_PDR13__pleiotropic drug resistance 13 |
| | | H1F | PSS | H1F_PSS | AT5G09710 | unspecified cations | Magnesium transporter CorA-like family protein |
| | | HD | PSS | HD_PSS | AT5G09710 | unspecified cations | Magnesium transporter CorA-like family protein |
| | | HD | PSS | HD_PSS | AT5G09930 | ABC transporters and multidrug resistance systems | ABCF2__ABC transporter family protein |
| | | DIAM | PVV | DIAM_PVV | AT5G59040 | metal | COPT3__copper transporter 3 |
| | | DIAM | SSV | DIAM_SSV | AT5G63060 | misc | Sec14p-like phosphatidylinositol transfer family protein |
| | | DIAM | PSV | DIAM_PSV | AT5G64410 | peptides and oligopeptides | ATOPT4_OPT4__oligopeptide transporter 4 |

**Table S8 Candidate genes related to one of the seven main categories listed by Subrahmaniam et *al*. 2018.**

| Categories (Subrahmaniam et *al.* 2018) | AGI | Class | Annotation |
|---|---|---|---|
| ABC transporters | AT3G28390 | transport | ABCB18_PGP18__P-glycoprotein 18 |
| ABC transporters | AT3G47740 | transport | ABCA3_ATATH2_ATH2__ABC2 homolog 2 |
| ABC transporters | AT4G15215 | transport | ABCG41_ATPDR13_PDR13__pleiotropic drug resistance 13 |
| ABC transporters | AT5G09930 | transport | ABCF2__ABC transporter family protein |
| cell wall modification and degradation | AT1G60390 | cell wall | PG1__polygalacturonase 1 |
| cell wall modification and degradation | AT1G70370 | cell wall | PG2__polygalacturonase 2 |
| cell wall modification and degradation | AT3G19620 | cell wall | Glycosyl hydrolase family protein |
| cell wall modification and degradation | AT3G53190 | cell wall | Pectin lyase-like superfamily protein |
| cell wall modification and degradation | AT3G56000 | cell wall | ATCSLA14_CSLA14__cellulose synthase like A14 |
| cell wall modification and degradation | AT4G15980 | cell wall | Plant invertase/pectin methylesterase inhibitor superfamily |
| cell wall modification and degradation | AT5G09760 | cell wall | Plant invertase/pectin methylesterase inhibitor superfamily |
| defense | AT1G69818 | stress | Defensin-like (DEFL) family protein |
| defense | AT4G12010 | stress | Disease resistance protein (TIR-NBS-LRR class) family |
| defense | AT5G63020 | stress | Disease resistance protein (CC-NBS-LRR class) family |
| histone modification | AT3G13170 | DNA | ATSPO11-1__Spo11/DNA topoisomerase VI, subunit A protein |
| histone modification | AT4G00660 | DNA | ATRH8_RH8__RNAhelicase-like 8 |
| histone modification | AT4G15570 | DNA | MAA3__P-loop containing nucleoside triphosphate hydrolases superfamily protein |
| histone modification | AT5G57160 | DNA | ATLIG4_LIG4__DNA ligase IV |
| histone modification | AT5G59970 | DNA | Histone superfamily protein |
| histone modification | AT5G09740 | RNA | HAC11_HAG05_HAG5_HAM2__histone acetyltransferase of the MYST family 2 |
| hormones | AT1G60380 | hormone metabolism | NAC (No Apical Meristem) domain transcriptional regulator superfamily protein |
| hormones | AT1G69700 | hormone metabolism | ATHVA22C_HVA22C__HVA22 homologue C |
| hormones | AT4G00880 | hormone metabolism | SAUR31__SAUR-like auxin-responsive protein family |

**Table S8 (Continued)**

| **Categories (Subrahmaniam et *al.* 2018)** | **AGI** | **Class** | **Annotation** |
|---|---|---|---|
| hormones | AT5G10720 | hormone metabolism | AHK5_CKI2_HK5__histidine kinase 5 |
| hormones | AT5G10990 | hormone metabolism | SAUR69__SAUR-like auxin-responsive protein family |
| hormones | AT5G25190 | hormone metabolism | ESE3__Integrase-type DNA-binding superfamily protein |
| meristem | AT1G23240 | development | Caleosin-related family protein |
| meristem | AT3G50670 | development | U1-70K_U1SNRNP__U1 small nuclear ribonucleoprotein-70K |
| meristem | AT4G00650 | development | FLA_FRI_RSB7__FRIGIDA-like protein |
| meristem | AT4G00800 | development | SETH5__transducin family protein / WD-40 repeat family protein |
| meristem | AT4G00930 | development | CIP4.1__COP1-interacting protein 4.1 |
| meristem | AT4G01430 | development | UMAMIT29__nodulin MtN21 /EamA-like transporter family protein |
| meristem | AT4G27410 | development | ANAC072_RD26__NAC (No Apical Meristem) domain transcriptional regulator superfamily protein |
| meristem | AT5G10830 | development | S-adenosyl-L-methionine-dependent methyltransferases superfamily protein |
| meristem | AT5G12330 | development | LRP1__Lateral root primordium (LRP) protein-related |
| photosynthesis | AT1G19150 | PS | LHCA2*1_Lhca6__photosystem I light harvesting complex gene 6 |
| photosynthesis | AT1G67740 | PS | PSBY_YCF32__photosystem II BY |
| photosynthesis | AT4G00895 | PS | ATPase, F1 complex, OSCP/delta subunit protein |
| photosynthesis | AT4G01330 | PS | Protein kinase superfamily protein |
